# PLK1 protects centromeres from BLM-mediated disintegration to promote chromosome biorientation

**DOI:** 10.1101/498808

**Authors:** Owen Addis Jones, Ankana Tiwari, Tomisin Olukoga, Alex Herbert, Kok-Lung Chan

**Affiliations:** Genome Damage and Stability Centre, University of Sussex, Brighton, BN1 7BG, United Kingdom

**Keywords:** PLK1, PICH/ERCC6L, BLM, ultrafine DNA bridge, PIT, chromosome biorientation, centromere dislocation, metaphase collapse

## Abstract

Faithful chromosome segregation cannot be achieved without proper chromosome biorientation (so-called metaphase alignment). Disabling PLK1, a key mitotic kinase, impairs multiple aspects of mitosis, particularly chromosome alignment. This is believed primarily due to unstable bipolar spindle-kinetochore attachments. Contrary to this belief, PLK1 inactivation does not necessarily abolish metaphase establishment, instead impairing its maintenance. Here, we demonstrate that the failure of chromosome biorientation maintenance is driven by a hitherto undescribed mechanism named ‘centromere dislocation’. Without an active PLK1 during mitosis, BLM helicase is illegitimately recruited to and unwinds a specific centromere domain underneath kinetochores in a PICH-dependent manner, impairing centromere configuration and rigidity. Acting with bipolar spindle pulling forces, the distorted centromeric chromatin is promptly converted into an ultra-thin DNA structure, termed ‘pre-anaphase interchromatin DNA threads’ (PITs) and fails to withstand spindle tension, resulting in whole-chromosome arms splitting. This devastating dechromatinisation action severely damages the centromere integrity and destroys normal metaphase maintenance. Therefore, our study unveils that in order to facilitate chromosome biorientation, PLK1 serves as a chromosome guardian to protect centromeres from disintegration, driven by a BLM-mediated chromatin decompaction activity.

---

Chromosome missegregation is widely implicated in cancers and birth defects^1^. In order to achieve faithful chromosome segregation, condensed chromosomes need to be correctly bi-oriented and aligned during metaphase. This requires the establishment of bipolar connections between spindle microtubules (MTs) that emanate from opposite centrosomes to each centromere via the macromolecular machinery of kinetochores (KTs)^2^. A single unattached chromosome can trigger the spindle assembly checkpoint (SAC), inhibiting the activation of the anaphase promoting complex/cyclosome (APC/C)^3^ and hence stop anaphase onset^4, 5^. This elegant surveillance system allows cells to correct KT-MT mis-attachment errors and prevent unlawful chromosome segregation. However, it inevitably keeps other bi-oriented chromosomes under prolonged spindle tension. Therefore, maintaining robust centromere structure and rigidity is crucial throughout mitosis but the underlying mechanism is not fully understood. Once the SAC is satisfied, the cleavage of cohesin, a ring-shaped complex that holds sister chromatids together occurs, triggering the disjunction of sister chromatids^6^. Intriguingly, the separated chromatids can remain intertwined by persistent DNA molecules that manifest as so-called ultrafine DNA bridges (UFBs)^7, 8^. It is believed that UFBs are resolved during anaphase by a UFB-binding complex, comprising of PICH (Plk1-interacting checkpoint helicase) translocase, BLM (Bloom’s syndrome) helicase and other interacting factors including TOP3A and TOP2^7–11^. However, its precise molecular action in resolving the intertwining chromatids remains largely unclear.

To facilitate mitotic progression, cells require a key regulatory kinase – Polo-like kinase 1 (PLK1). It has been shown that PLK1 plays a multifaceted role in mitosis, particularly during chromosome biorientation, which is proposed through stabilising KT-MT attachments^12–14^. Consistent with this notion, inhibition of PLK1 using the selective small molecule inhibitor, BI2536 (IC_50_ = 0.83nM)^15^, caused chromosome misalignment and induced a strong mitotic arrest in a way similar to the interferences of microtubule polymerisation using nocodazole, or of centrosome separation by monastrol (Supplementary Fig. 1a). However, by time-lapse microscopy on pre-synchronised normal diploid RPE1 cells (Supplementary Fig. 1b), we observed that unlike nocodazole and monastrol treatments, BI2536 did not completely abolish chromosome congression and metaphase establishment (Figs. 1a). The majority (~80%) of BI2536-treated RPE1 cells managed to align their chromosomes but shortly after succumbed to a loss of metaphase maintenance, namely chromosomes drifting away from the equator and dispersing into a ‘Fig-8’ or ‘polo’^16^ –like pattern (Figs. 1a-c, Supplementary Fig 1c and Movie S1). We referred to this post-metaphase misalignment phenomenon as ‘metaphase collapse’. In contrast, cells treated with the APC/C inhibitor, ProTAME, remained at metaphase for extended periods (Figs. 1a-c). More strikingly, we found that during ‘metaphase collapse’, cells produced a thread-like structure that was decorated with the PLK1 protein, which was not detected in untreated mitotic cells (Fig. 1d; arrows). Since the PLK1-coated threads were reminiscent of ultrafine DNA bridges (UFBs) that we and others identified previously in anaphase cells^7, 8^, we investigated whether they are DNA molecules; or alternatively a mis-localisation of PLK1 to cytoskeleton structures. We found that both well-known UFB-binding factors, PICH translocase and BLM helicase were present along the PLK1-coated thread structures (Figs. 1e, f and Supplementary Fig. 1d). Moreover, there was also a strong staining of replication protein A (RPA), indicating the presence of single-stranded DNA (Fig. 1f). The immunostaining results were further confirmed by examining Bloom’s syndrome fibroblast cells stably expressing a GFP-tagged BLM and RPE1 cells expressing a GFP-tagged PLK1 (Supplementary Figs. 1e, 1f; arrows). These data indicate that the PLK1-coated thread structures formed during ‘metaphase collapse’ are DNA molecules. Consistent with this, we also found that PLK1 is indeed a component of UFB-binding complex and localises to UFBs in normal anaphase cells (Supplementary Fig. 1g; arrows). As predicted, the DNA threads did not co-localise with mitotic microtubules (Supplementary Fig. 1h). Intriguingly, all DNA threads analysed (n=171) showed either one or both of their termini exclusively originating from centromeric regions (Fig. 1g). In some optical sections, it was obvious that two separating centromeres were interlinked by a DNA thread molecule (Fig. 1g; arrows). Alongside RPE1 cells, a short treatment of BI2536 in other normal human cells (including primary cells) and cancer cells also induced metaphase misalignment and centromeric DNA threads, albeit with different frequencies (Supplementary Fig. 2a; 82–6 (24%), 1BR3 (31%) and HCT116 (69%)). A plausible explanation for the formation of centromeric DNA threads could be due to precocious loss of sister chromatid cohesion, similar to the effect of SGO1 depletion^17, 18^, which leads to the exposure of sister DNA catenanes at centromeres^8^ (Supplementary Fig. 2b; arrows). However, consistent with published studies^19–21^, inhibition of PLK1 did not induce premature sister-chromatid separation in RPE1 cells (Supplementary Fig. 2c). Most crucially, by co-staining with the chromatid axis marker, TOP2A, we clearly visualised that chromosomes, which comprised of a pair of cohesed sister chromatids, were connected by DNA threads at their centromeres (Fig. 1h; arrows). Therefore, the loss of PLK1 function simultaneously induces collapse of metaphase and inter-chromosomal DNA entanglements. We hereby termed this new structure as <u>p</u>re-anaphase <u>i</u>nter-chromatin DNA <u>t</u>hread (PIT). Because of these striking findings, this led us to speculate that the loss of chromosome alignment may not be simply attributed to the unstable KT-MT attachment as previously thought.

**Figure 1.**
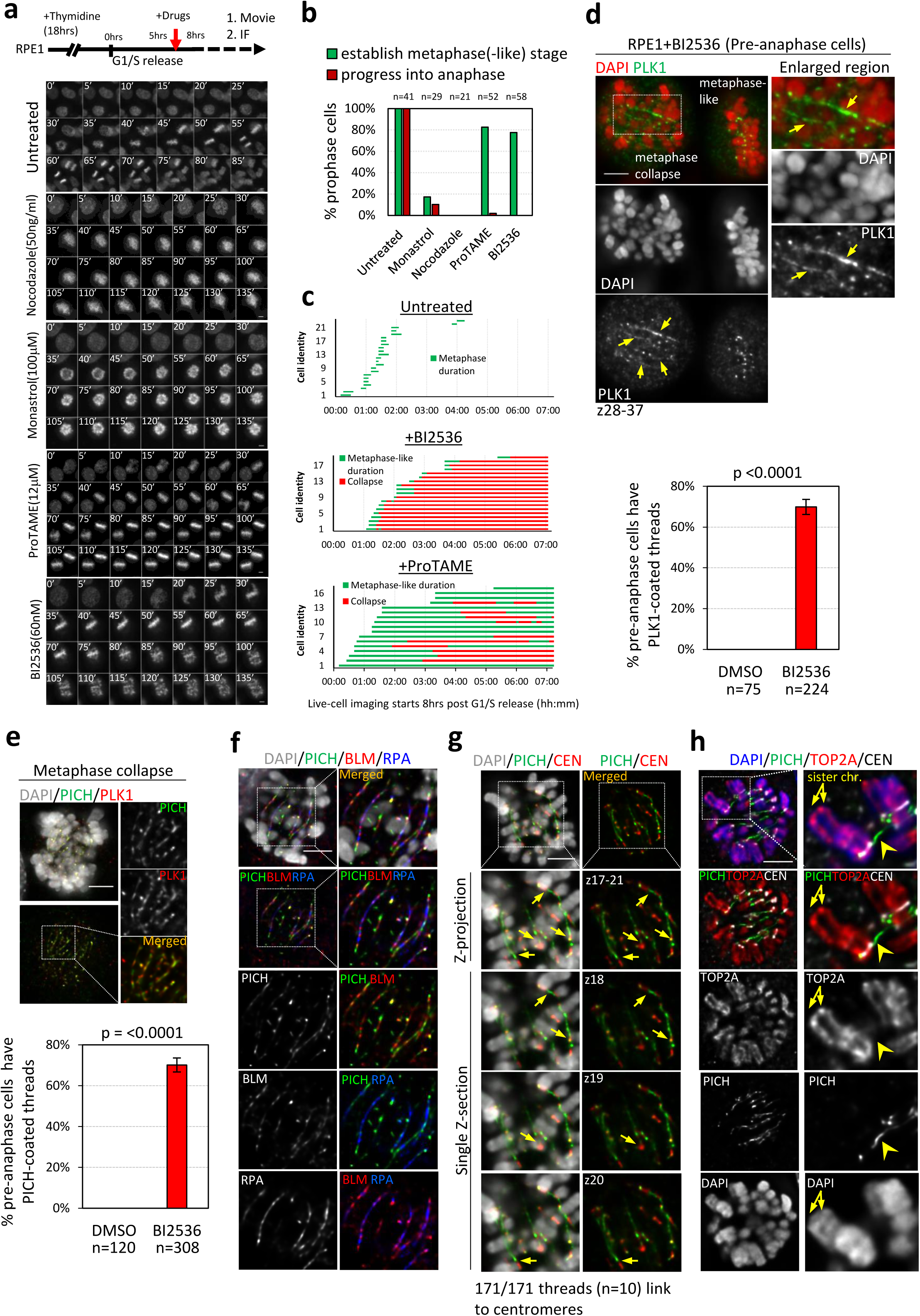
Plk1 inactivation induces inter-chromosomal DNA threads. **a**, Time-lapse imaging of mitotic progression in RPE1 cells treated with indicated inhibitors at 5hrs post thymidine-block and release. SiR-DNA was used to label DNA. **b**, Percentage of prophase cells progressing into metaphase(-like) and anaphase stages (n=number of prophase cells examined). **c**, Duration of metaphase(-like) stage in RPE1 cells treated with the indicated inhibitors. Note: only cells that reached metaphase(-like) stage were measured. **d**, PLK1 protein localises to thread structures (arrows) in the collapsed metaphase RPE1 cells after BI2536 treatment. Below: percentages of pre-anaphase cells (excluding prophase) showing PLK1-coated threads. **e**, PICH co-localises to PLK1-coated threads. Below: percentage of pre-anaphase cells (excluding prophase) showing PICH-coated threads **f**, Colocalisation of PICH, BLM and RPA on BI2536-induced DNA threads. Boxes indicate the enlarged regions. **g**, PICH-coated DNA threads linking centromeres (arrows). Below: panels showing consecutive images of single Z-planes with 200nm intervals. 171/171 (100%) threads were scored with centromere linkages. **h**, centromeric DNA threads (arrowhead) link chromosomes retaining a pair of sister chromatids (arrows), as labelled by TOP2A. Scale bar=5μm.

Given the highly repetitive nature of centromeres, we investigated whether PIT formation might be caused by inter-chromosomal homologous recombination that are unexpectedly induced by the PLK1 inhibition during (late) S phase. To address this, we used EdU to label on-going S-phase, but not G2 cells, while BI2536 was applied (Supplementary Fig. 3a). Contrary to our hypothesis, we found that the majority (69+4%) of the EdU-negative mitotic cells (which were in G2 phase) still generated PIT structures (Supplementary Fig. 3b), indicating that PIT formation is not caused by the interference of potential PLK1 functions during DNA replication. Moreover, treating early mitotic RPE1 cells (released 30mins after RO3306-induced G2 arrest) with BI2536 was also sufficient to induce metaphase collapse and PIT formation (Supplementary Fig. 3c). Therefore, the generation of the centromeric DNA interlinkage is a result of the impairment of PLK1’s mitotic function.

PICH translocase has been shown to possess a high binding affinity to DNA molecules under tension^22^, which may suggest that PIT formation may be a consequence of abnormal stretching of centromere chromatin. Interestingly, we detected activation of DNA damage responses as labelled by γH2AX staining at (peri)-centromere regions and that mainly occurred after the loss of metaphase alignment (Fig. 2a), implying that there is chromatin damage or remodeling during metaphase collapse. We next examined mitotic spread chromosomes obtained from pre-synchronised RPE1 cells after a short treatment of BI2536 (Supplementary Fig. 4a). As expected, in control (DMSO- and nocodazole-treated) RPE1 mitotic cells, their chromosomes displayed normal configurations and their average numbers were very close to 46 (diploid). In contrast, chromosomes from BI2536-treated cells became much shorter/compacted but strikingly, their chromosome numbers increased to an average of 59 (Supplementary Figs. 4b, c). This cannot be explained by chromosome mis-segregation because PLK1 inactivation blocks mitotic progression. Thus, this must be caused by chromosome fragmentation. Importantly, we found that the chromosome fragmentation is dependent on mitotic spindle tension as it was largely suppressed by co-treatment of the spindle poison, nocodazole (Supplementary Figs. 4b, c). By employing centromere-telomere fluorescence *in situ* hybridization (ctFISH), we detected that the fragmented chromosomes resembled telocentric chromosomes. However, in contrast their centromeric sides lacked telomeres (Fig. 2b; middle panels, asterisks), indicating that there is chromosomal breakage at/near the centromere. Supporting this, we observed partial separation of centromeres (Fig. 2b; middle panels, arrowheads) and occasionally the formation of an ultrafine centromeric DNA thread linking two separating chromosome arms (Fig. 2b; middle panels, connecting arrow). Combined with the cytological staining findings, these results strongly indicate that PIT formation is as a result of over-stretching of centromere chromatin by spindle pulling forces. As predicted, both centromere splitting and PIT formation were suppressed by nocodazole treatment (Fig. 2b and Supplementary Figs. 4d, e). To our knowledge, this form of spindle-dependent whole-chromosome arm rupture has never been described and we thus termed this as ‘centromere dislocation’. Using multi-color FISH (mFISH) karyotyping, we further proved that all fragmented chromosomes resulted from the splitting of the entire p-arm from the q-arm of normal chromosomes (Fig. 2c). In some cases, the separated whole-chromosome arms were located in close vicinity (Fig. 2c; e.g. chromosomes 7p-7q, 12p-12q, 17p-17q), which may imply a residual connection by the ultrafine centromeric DNA/PIT molecules. In addition, centromere dislocation tended to occur on longer chromosomes (Fig. 2c, inset). Collectively, our results demonstrate that in the absence of PLK1 function, the spindle pulling force can cause a longitudinal stretching of the centromere axes, resulting in whole chromosome arm separation and PIT formation (Fig. 2d). Indeed, we were able to detect a chromosome axial element, condensin, associating along the PIT structures (Fig. 2e; arrows). Therefore, these findings provide a new explanation of how chromosome misalignment may arise, namely the loss of centromere integrity.

**Figure 2.**
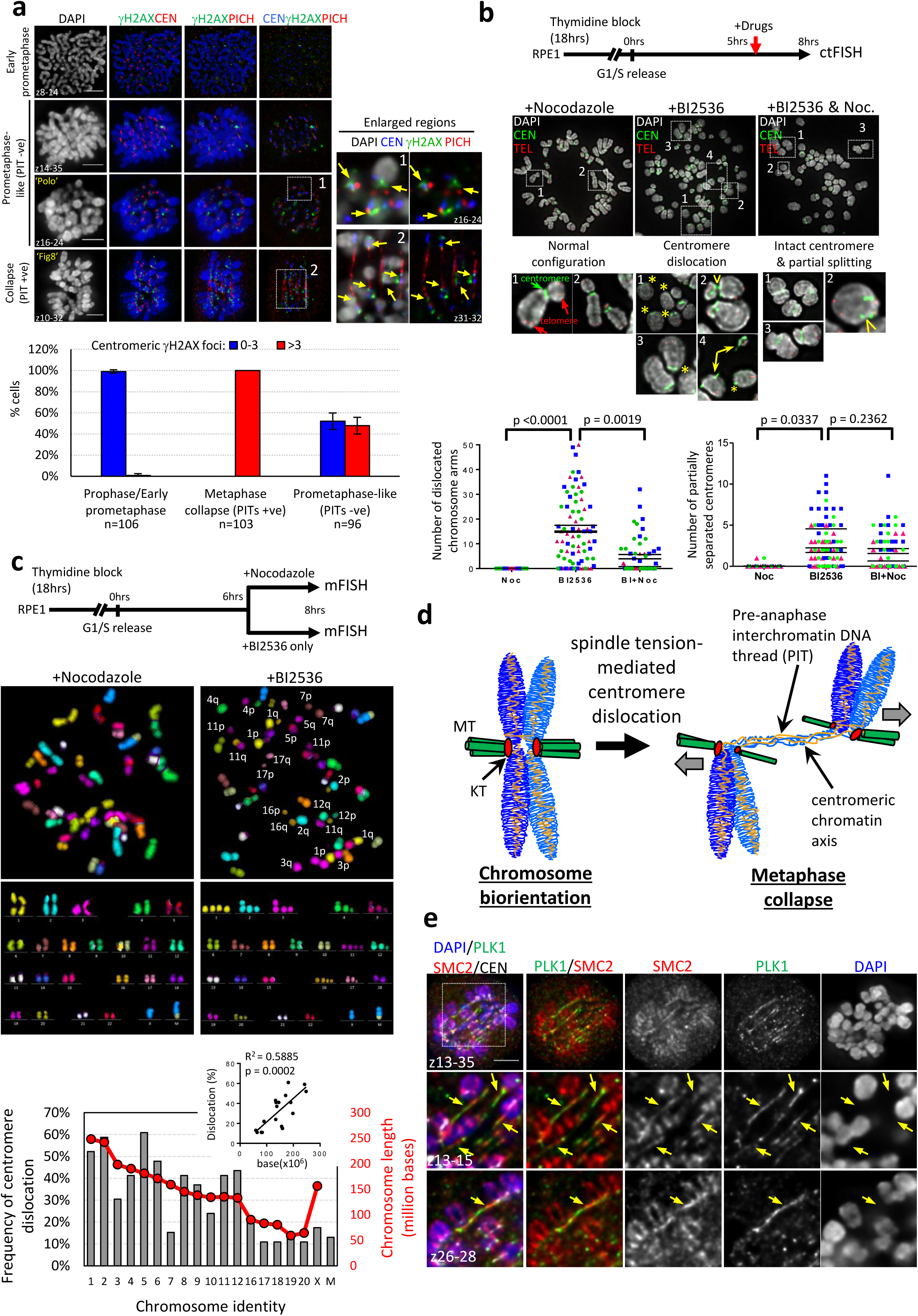
Centromere dislocation leads to PIT formation. **a**, DNA damage responses occur during the formation of PITs. Deconvolved images showing different mitotic phases of RPE1 cells after BI2536 treatment stained with PICH, γH2AX and CEN antibodies. Enlarged boxes: arrows denote γH2AX signals at centromeric regions positive for PICH-coated PITs, or strong PICH foci. Below: quantitation of mitotic populations showing centromeric γH2AX foci (n=total number of cells scored in three independent experiments). **b**, BI2536 treatment induces spindle-dependent centromere dislocation. Images of chromosome spreads from pre-synchronised RPE1 cells treated with indicated inhibitors. Below: enlarged panels showing chromosomes with intact configurations (left), centromere dislocation (middle; asterisks) and partial separation of centromere (middle & right; arrowheads). A yellow arrow denotes an ultra-thin centromeric DNA thread linking two splitting arms (middle#4). Bottom; quantitation of the number of complete (left) and partial (right) splitting whole-chromosome arms per spread (75 spreads from three independent experiments were analysed and means from each experiment are shown). **c**, mFISH karyotypes showing whole-chromosome arm splitting. RPE1 cells were treated with nocodazole or BI2536 alone at 6hrs post G1/S release. Note: There is a marker ‘M’ chromosome with a translocation of chromosome X and 10 in RPE1 cells. Bottom; frequencies of centromere dislocation among the indicated chromosomes. Inset showing positive correlation between chromosome length and centromere dislocation (23 spreads were analysed. Note: acrocentric chromosomes were not determined and the length of ‘marker’ chromosome is unknown). **d**, A diagram depicts the formation of PITs during centromere dislocation in a spindle tension-dependent manner. **e**, The chromosmal axial element, condensin SMC2, associates along PITs (arrows). Scale bar=5μm.

To rule out any off-target effects of the BI2536 treatment, we repeated experiments using an engineered RPE1 cell line where the endogenous wildtype PLK1 has been replaced with an analog-sensitive allele (PLK1as), where its catalytic cavity has been modified not to be bound by BI2536 inhibitor but instead to an ATP-analog, 3-MB-PP1^23^. As predicted, BI2536 treatment failed to induce both metaphase collapse and PIT formation in PLK1as RPE1 cells. However, these mitotic defects were phenocopied by using the 3-MB-PP1 analog (Supplementary Figs. 5a-d). In addition, PLK1 depletion through RNAi also induced PIT formation and centromere dislocation, ruling out potential dominant effects of trapping an inactive form of PLK1 onto chromatin by either BI2536 or 3-MB-PP1 (Supplementary Figs. 5e, f). Thus, mitotic PLK1 kinase activity *per se* is crucial for the maintenance of centromere rigidity.

The failure to withstand bipolar spindle pulling may suggest that centromere structures are improperly established in the absence of mitotic PLK1 activity. Careful analyses of different mitotic phases of BI2536-treated cells revealed that prior to metaphase collapse there is a progressive accumulation of RPA foci at/near kinetochores, from prometaphase to metaphase-(like) cells (Fig. 3a), indicating increased formation of ssDNA structures at centromeres, which again was dependent on spindle pulling forces (Fig. 3a). A recent study has reported that phospho-RPA foci are observed at centromeres on cytospun metaphase chromosomes^24^. However, we rarely detected RPA foci at centromeres in untreated intact metaphase cells (Fig. 3b). Thus, our data infer that in the absence of PLK1 activity, chromosome congression mediated by spindle pulling may trigger DNA melting at the centromere. However, we also detected a striking accumulation of BLM helicase at/near kinetochores in the metaphase(-like) cells, again only after BI2536 treatment (Figs 3c). In addition, strong PICH foci were observed at kinetochores^7^ and occasionally at the core centre of some centromeres (Supplementary Fig. 6a; yellow arrows). The latter presumably represents stretched DNA catenation structures in between sister centromeres, as previously proposed^7, 8, 10, 25, 26^. PICH has been reported to localise at kinetochores in normal mitotic cells^7^. To confirm that PLK1 inactivation not only leads to BLM but also PICH over-loading at kinetochores, we employed quantitative imaging on co-cultured cells containing both wildtype RPE1, together with the modified RPE1 cells that express GFP-tagged PLK1as protein. This allowed us to directly compare the focal intensities of PICH. We consistently detected significant elevations of both number and intensities of PICH foci at kinetochores in the parental RPE1 cells compared to the GFP-PLK1as RPE1 cells, following a short treatment of BI2536 (Supplementary Figs. 6b-d). Conversely, 3-MB-PP1 treatment led to strong PICH accumulation in the PLK1as RPE1 cells (Supplementary Figs. 6b-d). As predicted, PICH, BLM and RPA foci mostly colocalised at the same kinetochores (Supplementary Fig. 6e; arrows). Given that BLM helicase and PICH translocase have strong activities to unwind or displace DNA molecules^22, 27^, we speculated that, apart from the possibility of microtubule pulling forces, the ssDNA formation at centromeres may be driven by the BLM and/or PICH protein, which subsequently weakens the centromere rigidity.

**Figure 3.**
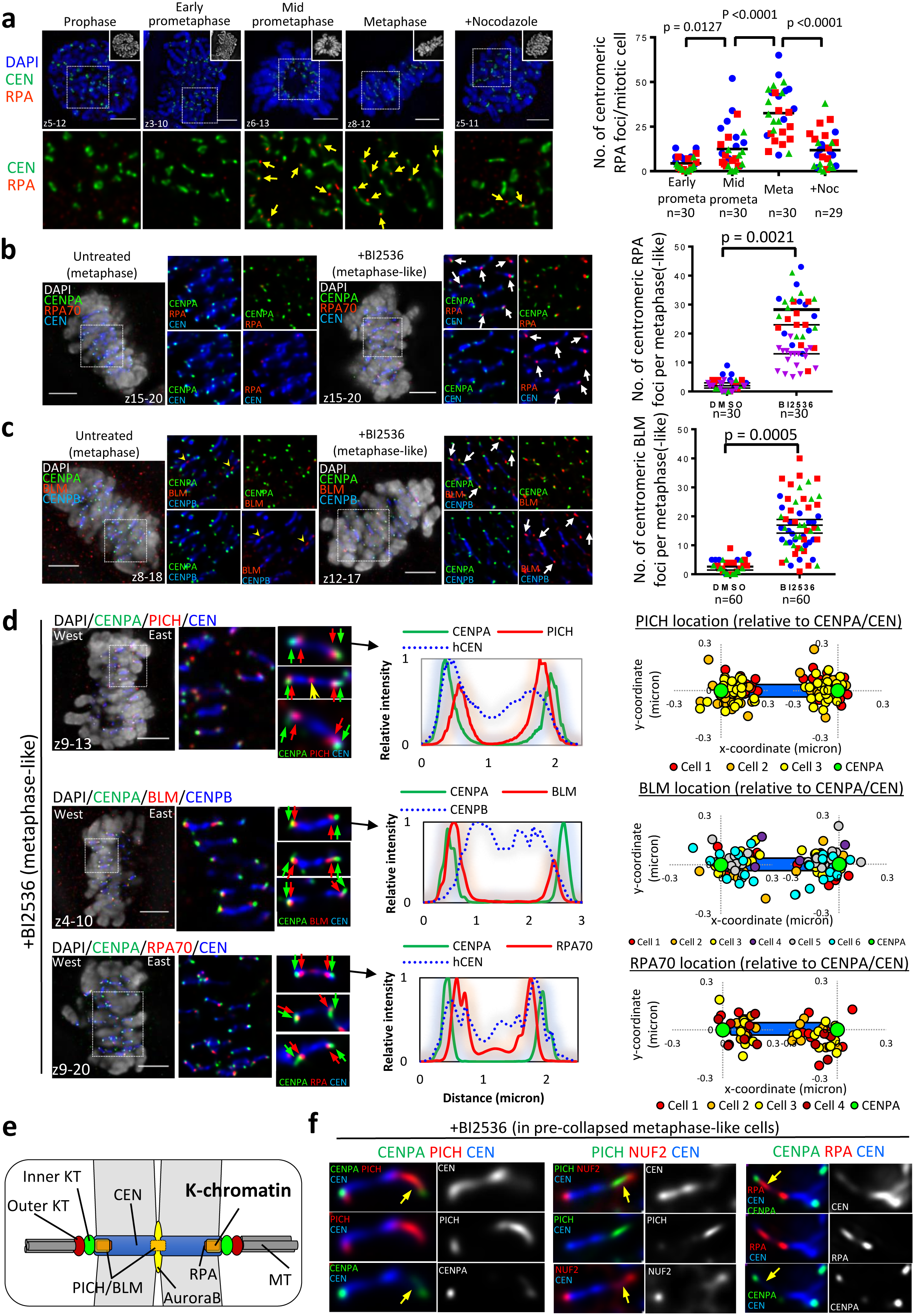
Illegitimate ssDNA formation at centromere K-chromatin sites. **a**, Plk1 inhibition induces the progressive formation of RPA foci at kinetochores from early prometaphase to metaphase. Right: quantification of RPA foci during each mitotic stage. **b**, Examples of metaphase(-like) RPE1 cells showing RPA foci at/near kinetochores (arrows) following BI2536 treatment. Right: quantitation of centromeric RPA foci per metaphase(-like) cell. **c**, same as (b) but analysis of centromeric BLM foci (n=total number of cells scored from 3 independent experiments; mean from each experiment is shown). **d**, Representative images showing the relative locations of PICH, BLM and RPA70 foci compared to CENPA and core centromeres (CEN) in BI2536-treated metaphase-like cells. Middle panels: scanline measurements of the relative positions of different proteins within centromeres. Right panels: location coordinates of PICH, BLM and RPA foci relative to CENPA foci and central centromere on the opposite sides of sister KT. **e**, A diagram depicts the K-chromatin site underneath kinetochores bound by UFB-binding factors. **f**, Examples of kinetochore detachment from core centromeres after BI2536 treatment in metaphase(-like) RPE1 cells. Arrows indicating the detached KTs remain linked by a short DNA thread as labelled by UFB-binding proteins. Scale bar=5μm. All RPE1 cells were pre-synchronised at G1/S by single thymidine block, followed by drug treatments at 5hrs post release. Immunostaining was performed at 8hrs post release. Scale bar=5μm.

Further support to this idea came from our high-precision localisation mapping of PICH, BLM and RPA foci at kinetochores/centromeres (Supplementary Figs. 7a-c). Although PICH translocase has been reported to co-localise with PLK1 at kinetochore regions, its localisation is independent of PLK1^10, 28^ (Supplementary Fig. 6e). Surprisingly, we found that PICH indeed did not actually locate within kinetochore complexes as previously thought, instead it was found at a position towards the centromere chromatin, approximately 160nm away from the outer kinetochore component, NUF2 (Supplementary Fig. 7d, f). As a control comparison the inner KT component, CENPA, was mapped only ~100nm inwards from NUF2 in metaphase cells, and the distance reduced to ~80nm in anaphase cells (Supplementary Figs. 7e, f), probably because of a reduction of intra-kinetochore tension after sister-chromatid cohesion loss^29, 30^. Nevertheless, further co-staining of PICH and CENPA confirmed that PICH localises separately from the inner kinetochore and resides at a centromeric DNA domain ~100nm beneath CENPA (Figs. 3d and Supplementary Fig. 7f). Likewise, both BLM and RPA foci, were also mapped underneath the inner kinetochore, as labelled by CENPA, with distances of ~120 and ~150nm, respectively, which are closely adjacent to PICH (Figs. 3d and Supplementary Fig. 7f). All of these proteins displayed mirror localization patterns between opposite sister kinetochores, reflecting a true biological relevance of centromere symmetry. To distinguish this specific DNA domain of the centromere, we thereby referred to it as ‘kinetochore-chromatin or K-chromatin’ (Fig. 3e). These new cytological localisation findings support our notion that centromeric DNA is specifically targeted by the UFB-binding proteins, PICH and BLM that may cause the centromere fragility under spindle pulling forces. Indeed, we detected detachment of kinetochores from core centromeres in pre-collapsed metaphase cells (Fig. 3f and Supplementary Fig. 8a; arrows). We believe this represents the early stage of centromere deformation. Moreover, as predicted from our model that the bipolar spindle pulling induces longitudinal separation of centromeres (Fig. 2d), we detected a large percentage of the centromeres losing one kinetochore after metaphase collapse (Supplementary Figs. 8b-d). Notably, the centromeric sides missing kinetochores were concomitant with the formation of a PIT molecule (Supplementary Fig. 8c; connecting arrows). Therefore, prior to metaphase collapse, centromeres are undergoing apparent deformation, which may predispose to the subsequent centromere dislocation (Supplementary Fig. 8e and see model in Fig. 6).

To further prove that BLM and/or the PICH complex initialises centromere deformation, we knocked down BLM in synchronised RPE1 cells by RNAi. Silencing BLM longer than 48 hours reduced the efficiency of thymidine release, therefore we treated cells with siBLM oligo for only 24 hours prior to G1/S release. Despite the partial depletion (Supplementary Figs. 9a-c), BLM knockdown remarkably reduced RPA foci formation at the K-chromatin region induced by BI2536 in the metaphase(-like) populations (Figs. 4a). This strongly indicates that the formation of centromeric ssDNA is dependent on BLM. More crucially, knocking down BLM also significantly diminished both the formation of PITs and centromere dislocations, as compared to control siRNA transfection (Supplementary Figs. 9d, e). However, BLM depletion did not impair PICH’s centromeric accumulation (Fig. 4a). Thus, PICH alone is not sufficient to drive centromere dislocation. To confirm the specificity of BLM knockdown, we performed our analyses on HAP1 cells in which the endogenous BLM was knocked out by CRISPR genome editing^31^ (Supplementary Fig. 9f). Consistent to the RNAi results, BLM knockout in HAP1 cells again abolished PIT formation and centromere dislocation (Fig. 4b, c). Although we occasionally detected chromatid breaks in ΔBLM HAP1 cells, their breakpoints were not at centromeres (Fig. 4c, arrow). In agreement with our hypothesis, BLM’s unwinding activity is essential in order to trigger centromere disintegration, as a BLM mutant (Q672R) lacking ATPase and helicase activity was unable to induce both PIT formation and centromere dislocation, despite its accumulation at the K-chromatin region of centromeres (Figs. 4d, e and Supplementary Fig. 9g). Interestingly, although PICH alone is incapable of driving centromere disintegration, its depletion on the other hand could suppress centromeric ssDNA formation, PITs and centromere dislocations (Figs. 4f, g and Supplementary Fig. 9h). We found that this is because PICH acts upstream in the recruitment of BLM to centromeres (Fig. 4h). Therefore, our results demonstrate that the main cause of centromere disintegration is driven by illegitimate unwinding of centromeric DNA by the BLM helicase, which destroys normal centromere architectures and rigidity, leading to the failure to withstand spindle pulling forces.

**Figure 4.**
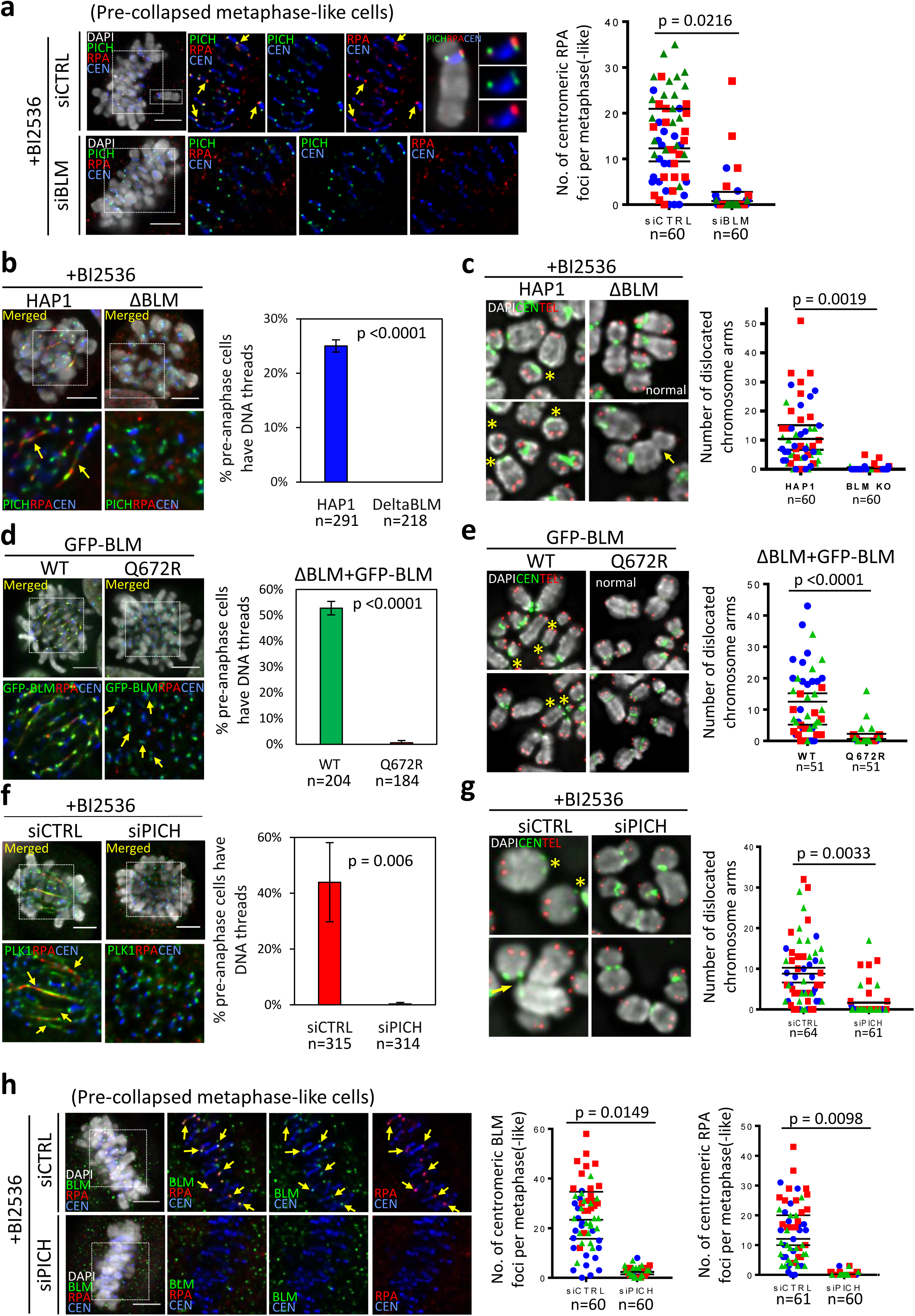
BLM helicase drives the formation of centromeric ssDNA, PITs and centromere dislocation. **a**, BLM knockdown diminished centromeric RPA but not PICH foci in metaphase(-like) RPE1 cells. Left: representative images of PICH and RPA centromeric foci in siCtrl and siBLM RPE1 cells. Right: quantitation of centromeric RPA foci per cell (mean from three independent experiments shown). **b**, Quantification of PITs in BLM knockout HAP1 cells shows an absence of PIT formation (mean ± S.D. from three independent experiments). **c**, Quantification of centromere dislocation in BLM knockout HAP1 cells (asterisk, chromosome dislocations; arrow, chromatid breaks) (mean from three independent experiments shown). **d**, Quantitation of PIT formation in ΔBLM HAP1 cells stably expressing a GPF-tagged wildtype BLM or Q672R mutant (mean ± S.D. from three independent experiments). **e**, Similar to (d) but quantitation of centromere dislocation (mean from three independent experiments shown). **f**, PICH knockdown in RPE1 cells abolished PIT formation (mean ± S.D. from three independent experiments). **g**, PICH knockdown in RPE1 cells diminished centromere dislocation (asterisk, chromosome dislocations; arrow, partial dislocation) (mean from three independent experiments shown). **h**, Loss of both BLM and RPA foci at centromeres in siPICH RPE1 metaphase-like cells (mean from three independent experiments shown). All analyses were done in cells released from single-thymidine block following a short treatment of BI2536. Scale bar=5μm. n=total numbers of cells analysed.

Our data, thus far, strongly argue that a crucial function of PLK1 during chromosome biorientation is to protect centromere integrity rather than merely securing stable KT-MT attachments. To further test this, we examined the effect of PLK1 inactivation in mitotic cells only after they had fully established chromosome biorientation at metaphase. RPE1 cells stably expressing a GFP-tagged PLK1 protein were blocked at metaphase using the APC/C inhibitor, ProTAME. High-resolution live-cell imaging recorded that upon the addition of BI2536, the aligned metaphase chromosomes started losing chromosome biorientation as predicted. However, most importantly, this was accompanied with the formation of PITs (Fig 5a and Movies S2, S3), indicating the occurrence of centromere dislocation. Furthermore, we also found that centromere dislocation occurs rapidly, as more than 60% of the metaphase-arrest cells exhibited misaligned chromosomes with PIT formation after just 30 minutes of PLK1 inhibition (Fig. 5b). Since PIT formation is dependent on spindle tension, these data thus imply that the chromosome misalignment cannot be due to the loss of KT-MT connection. Supporting this, depleting PICH or BLM, which has been shown to abolish centromere dislocation, significantly prolonged metaphase maintenance following PLK1 inhibition (Fig. 5c and Movies S4-S6). Therefore, the loss of chromosome alignment is primarily triggered by BLM-mediated centromere disintegration rather than KT-MT destabilisation. It is worth to note that chromosome misalignment inevitably occurred in the PICH- and BLM-depleted cells after significant delays, but their distinct ‘polo’ misalignment pattern suggests this is likely due to KT-MT detachment. These results, therefore, also imply that the spindle destabilisation is a relatively slow process. Taken together, we propose a novel role of PLK1 in promoting chromosome biorientation, which is to protect centromeres from BLM-mediated dechromatinisation under bipolar spindle tension (Fig. 6).

**Figure 5.**
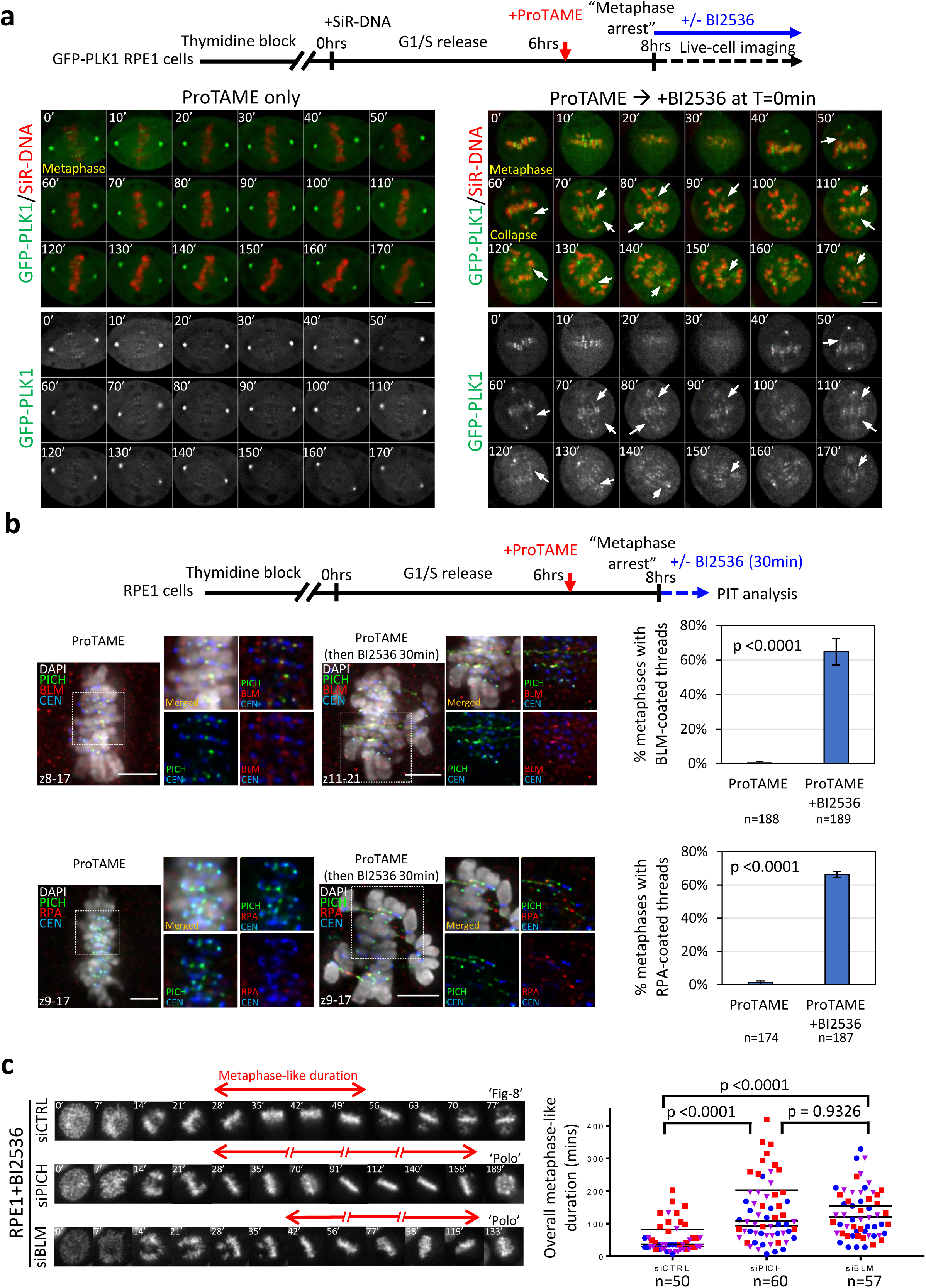
PICH and BLM depletion suppresses metaphase collapses caused by centromere disintegration. **a**, PLK1 inactivation after metaphase establishment triggers PIT formation and metaphase collapse. Live-cell imaging was carried out on GFP-tagged PLK1 RPE1 cells arrested in metaphase by the APC/C inhibitor, ProTAME. The positions of metaphase cells were recorded and live-cell imaging started immediately after the addition of 60nM of BI2536 (Time=0). Arrows indicate the formation of DNA threads coated by GFP-PLK1 protein. **b**, RPE1 cells arrested at metaphases by ProTAME treatments lost the maintenance of chromosome alignment and generated PITs within 30min after the addition of BI2536. PITs are labelled by PICH, BLM and RPA staining (mean ± S.D. from three independent experiments are shown). Scale bar=5μm. **c**, Depletion of PICH and BLM prolongs metaphase-(like) stage of RPE1 cells after PLK1 inhibition. Time-lapse microscopy of mitotic progression in the indicated siRNA oligos transfected into RPE1 cells under BI2536 treatment (Black bars indicate the metaphase-like stages). Overall duration of metaphase(-like) stage in control, PICH- and BLM-depleted RPE1 cells following BI2536 treatment (mean from three independent experiments shown). n=total number of cells scored.

**Figure 6.**
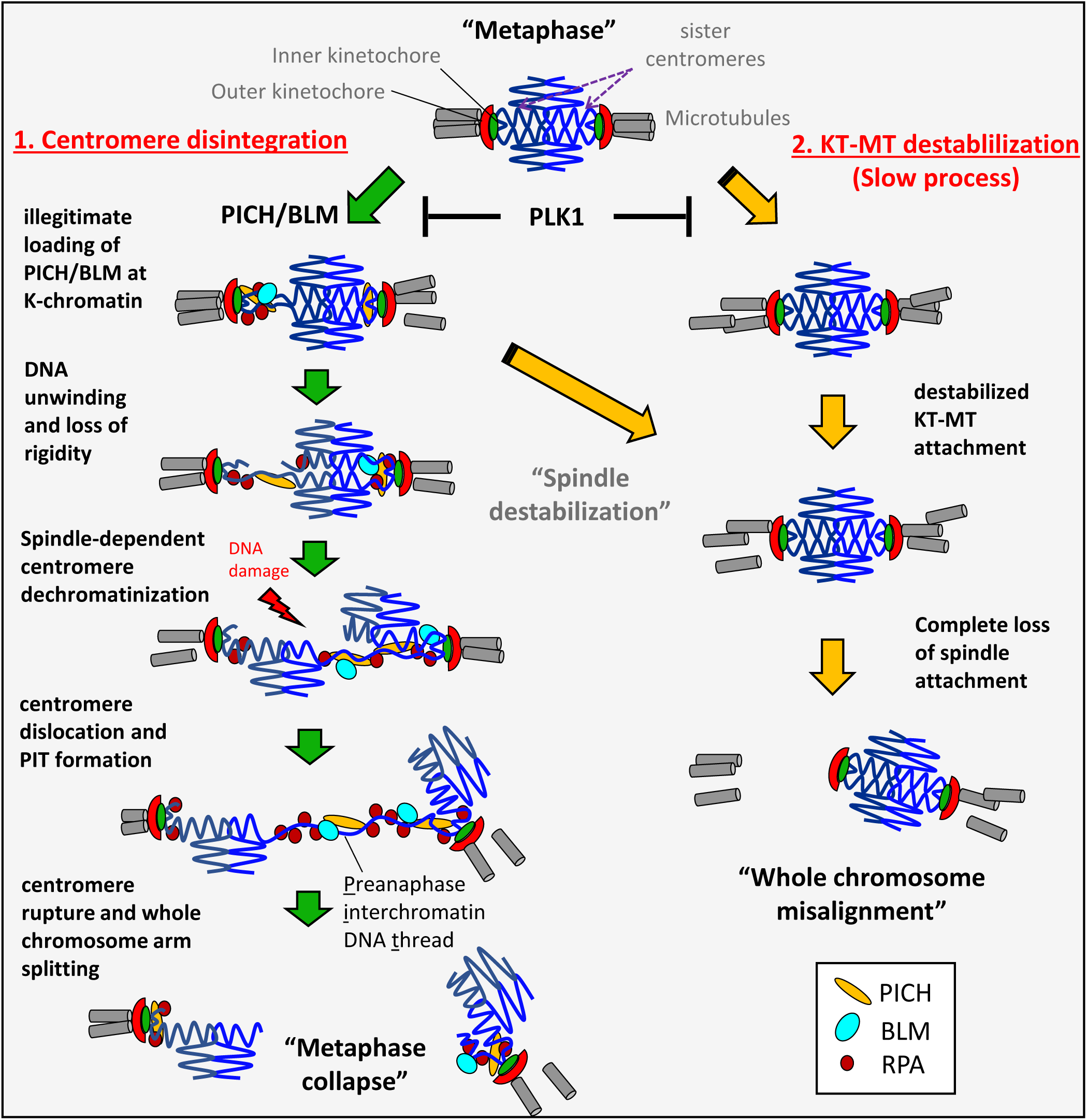
A model of chromosome misalignment driven by BLM/spindle-dependent centromere disintegration. PLK1 promotes the maintenance of chromosome biorientation and metaphase alignment through two important roles. First, it protects centromeres from disintegration, which is triggered by a PICH/BLM-mediated centromeric DNA unwinding followed by spindle-dependent chromatin decompaction. This results in centromere rupture, whole-chromosome arms splitting and metaphase collapse. Second, PLK1 maintains stable kinetochore-microtubule attachment. However, once the KT-MT connection is established, the loss of PLK1 activity does not cause an immediate KT-MT destabilisation and whole chromosome misalignment, which undergoes a relatively slow process and the spindle detachment may also simultaneously result from the loss of centromere integrity.

Chromosome alignment failure is generally attributed to the disorganisation and/or assembly errors of spindle-kinetochore apparatus. However, our current study provides a novel insight into how chromatin remodeling and repair factors have a huge influence on proper chromosome biorientation, and also highlights the importance of the protection of centromere integrity mediated by PLK1. It is reasonable to think that centromere disintegration may be caused by defects in chromatin condensation, as it has been shown that depletion of condensin I leads to abnormally stretched centromere structures and increased inter-kinetochore distances^32^. However, the phenotypes reported here are more distinct and severe. Crucially, we demonstrate that centromere dislocation and rupture is not a passive process simply mediated by spindle tension. In fact, it is actively driven through the unwinding of centromeric DNA by the PICH/BLM complex. Importantly, the identification of this centromere-specific breakage pathway, independent of the necessity of passing through chromosome mis-segregation^33, 34^, also provides us an alternative explanation on how whole-chromosome arm rearrangements may initiate, which are observed in many human tumours and rare genetic disorders^35–37^.

Another very intriguing finding is that the PICH/BLM complex is capable of triggering chromatin decompaction when it acts in concert with bipolar spindle pulling forces. Within a short period of PLK1 inhibition, the bulky centromere chromatin is converted into an ultra-thin DNA thread structure that is reminiscent of UFBs observed in anaphase cells^7, 8^. It has been suggested that the association of PICH/BLM complex on DNA bridging structures facilitates their resolution, however, its precise molecular action is yet to be elucidated. Our current finding provides an alternative proposal that the biochemical and biophysical co-action exerted by PICH/BLM-mediated DNA unwinding and spindle pulling may introduce localised dechromatinisation of the intertwined sister chromatids that relaxes the entanglement constraints to facilitate their timely segregation. Thus, it is conceivable that such powerful and potent dechromatinisation activity requires tight regulation, possibly via phosphorylation by mitotic kinases such as PLK1 in order to prevent unwanted pathological damage on mitotic chromatin, as reported here. In fact, both PICH and BLM are substrates of mitotic kinases, including PLK1^7, 38, 39^ and it has been shown that hyper-phosphorylated BLM exhibits poor association to mitotic chromatin and on sister DNA bridges, induced by Sgo1-depletion^8, 39, 40^. However, there is still a possibility that PLK1 may act simultaneously to strengthen centromere structure and to suppress DNA remodeling activities. In conclusion, our study unveils an unexpected interplay between PLK1 and DNA remodeling/repair factors in safeguarding centromere integrity to facilitate proper chromosome biorientation.

## ACKNOWLEDGEMENTS

We would like to thank the people in the Genome Centre for their great support. We thank Robert Lera and Mark Burkard for providing us with the RPE1 derivative cells, Marcel van Vugt for HAP1 and HAP1 BLM knockout cells, and Phillip North for GM08505 BS cells. We thank Jessica Hudson for BLM siRNA oligos. We also thank Kim Nasmyth, Jon Baxter and Mark Burkard for helpful comments on the manuscript. This work is supported by Sir Henry Dale Fellowship (Ref: 104178/Z/14/Z) from Wellcome Trust and the Royal Society, and by the Genome Damage and Stability Centre. K.L.C. is the recipient of Sir Henry Dale Fellowship.

## AUTHOR CONTRIBUTIONS

O.AJ., A.T. and K.L.C designed and performed the experiments with help from T.O. and A.H. K.L.C wrote the manuscript with input from all authors.

## METHODS

### Cell culture

RPE1-hTERT, 82–6-hTERT normal diploid cell lines, 1BR3 primary fibroblasts, HCT116 colon cancer cells were obtained from the Genome Damage and Stability Centre Cell Bank. RPE1-hTERT derivative cells were generated and supplied by Mark Burkard (University of Wisconsin). Bloom’s syndrome fibroblasts (GM08505) were obtained from Phillip North (University of Oxford). HAP1 cells and HAP1 *Δ*BLM cells were obtained from Marcel van Vugt (University of Groningen). All cell lines passed mycoplasma tests (Lonza MycoAlert kit). RPE1-hTERT and its derivative cells were grown in DMEM/F-12 medium (Sigma) containing 15% foetal calf serum (FCS) and Pen/Strep antibiotics (P/S). 82–6 fibroblast cells were grown in DMEM/F-12 medium containing 15% FCS and P/S. HAP1 cells were grown in IMDM (Gibco) containing 10% FCS and P/S. 1BR3 primary cells were grown in MEM (Gibco) containing 2mM L-glutamine, 15% FCS and P/S. HCT116 cells were grown in McCoy’s 5A (Gibco) containing 15% FCS and P/S. Bloom’s syndrome fibroblasts (GM08505) were transfected with a pEGFP-hBLM construct and selected by 700μg/ml G418 for 14 days. A single clone was isolated and maintained in MEM (Gibco) containing 2mM L-glutamine, 10% FCS, P/S and G418. Cell cultures were maintained at 37°C in a humidified atmosphere containing 5% CO2. GFP-BLM(WT) and GFP-BLM(Q672R) HAP1 cells were generated by stable transfection with the corresponding constructs in HAP1 *Δ*BLM cells by using FuGene HD (Promega) according to the manufacturer’s guidelines. The DNA constructs were created by sub-cloning EGFP-hBLM (WT) or Q672R (helicase dead mutant) fragments into a pSYC-181-(NEO) vector. Following a 1.2mg/ml of G418 selection for 14 days, GFP positive populations were sorted and isolated using a FACS cell sorter (BD FACSMellody).

### Cell synchronisation and drug treatments for mitotic cell analysis

Cells were treated with 2mM of thymidine for 18 hours to enrich cells at the G1/S boundary. Cells were then released into S-phase by washing three times with pre-warmed culturing medium, or pre-warmed 1xPBS and released into fresh medium. Five to six hours post G1/S release, indicated inhibitors were added. At approximately 8–9 hours post the G1/S release, mitotic cells were fixed or enriched for analyses.

### RNA interference

Cells were transfected with siRNA oligonucleotides using Lipofectamine RNAiMAX transfection reagent (Thermo Fisher Scientific) following the manufacturer’s guidelines. Cells underwent 1 or 2 rounds of siRNA transfection as necessary.

Non targeting pool (Dharmacon ON-TARGET plus Non-targeting Pool – D-001810–10–05. UGGUUUACAUGUCGACUAA; UGGUUUACAUGUUGUGUGA; UGGUUUACAUGUUUUCUGA; UGGUUUACAUGUUUUCCUA) PLK1 siRNA sequence (Dharmacon ON-TARGET plus SMARTpool – L-003290-00-0005. GCACAUACCGCCUGAGUCU; CCACCAAGGUUUUCGAUUG; GCUCUUCAAUGACUCAACA; UCUCAAGGCCUCCUAAUAG) Sgo1 siRNA sequence (Dharmacon ON-TARGET plus SMARTpool – L-015475-00-0005. CAGCCAGCGUGAACUAUAA; GUUACUAUCUCACAUGUCA; AAACGCAGGUCUUUUAUAG; GUGAAGGAUUUACCGCAAA) BLM siRNA sequence (Dharmacon ON-TARGET plus Individual – J-007287-08-0005. GGAUGACUCAGAAUGGUUA) PICH siRNA sequence (Invitrogen - AAUUCGGUAAACUCUAUCCACAGCU)

### Fluorescence immunostaining

For immunostaining analyses, cells were seeded onto No. 1.5 or No. 1.5H cover glass and fixed with Triton X-100-PFA buffer (250 mM HEPES, 1xPBS, pH7.4, 0.1% Triton X-100, 4% methanol-free paraformaldehyde) at 4°C for 20mins, or with PBS-PFA buffer (1xPBS, 4% methanol-free paraformaldehyde) at room temperature for 10mins. Pre-extraction was carried out in indicated experiments before fixation by incubation of the cover glass in pre-extraction buffer (20mM HEPES pH7.4, 0.5% Triton X-100, 50mM NaCl, 3mM MgCl_2_, 300mM sucrose) for 10–15secs.

Primary antibodies used: anti-PICH (Abnova; H00054821-B01P, 1:100), anti-PICH (Abnova; H00054821-D01P, 1:100), anti-BLM (Santa Cruz; sc-7790, 1:50), anti-BLM (Abcam; ab2179, 1:200), anti-γH2AX (Upstate; JBW-301, 1:400), anti-TOP2A (Santa Cruz; sc-5348, 1:100), anti-SMC2 (Bethyl Lab; A300–058A, 1:200), anti-RPA70 (Abcam; ab79398, 1:200), anti-RPA32 (Abcam; ab2175, 1:200), anti-CENPA (Abcam; ab13939, 1:100), anti-CENPB (Abcam; ab25734, 1:800), anti-NUF2 (Abcam; ab122962, 1:200), anti-PLK1 (Santa Cruz; sc-55504, 1:100), anti-pericentrin (Abcam; ab4448, 1:400), anti-centromere (ImmunoVision; HCT-0100, 1:400) and GFP booster (ChromoTek; gba-488, 1:200). Secondary antibodies used: donkey anti-mouse Alexa Fluor 488, 555 and 647; donkey anti-rabbit Alexa Fluor 488, 555 and 647; donkey anti-goat Alexa Fluor 488 and 555; goat antihuman DyLight 550 and 650 (All secondary antibodies used at 1:500 dilution). Immunofluorescence staining was performed according to previously described protocols^8^. Cells were mounted using Vectashield containing DAPI.

### High-resolution deconvolution microscopy

Images were acquired under a Zeiss AxioObserver Z1 epifluorescence microscopy system with 40x/1.3 oil Plan-Apochromat, 63x/1.4 oil Plan-Aprochromat and 100x/1.4 oil Plan-Aprochromat objectives and a Hamamatsu ORCA-Flash4.0 LT Plus camera. The system is calibrated and aligned by using 200nm-diameter TetraSpeck microspheres (ThermoFisher). Ten to fifty z-stacking images were acquired at 200nm intervals covering a range from 2–10μm by using ZEN Blue software.

Deconvolution was carried out using Huygens Professional deconvolution software (SVI) with a measured point-spread-function (PSF) generated by 200nm-diameter TetraSpeck microspheres. Classical maximum likelihood estimation method with iterations of 40 to 60 and signal-to-noise of 20 to 60 was applied.

### Time-lapse Live-cell microscopy

Cells were seeded on 2-well or 4-well tissue culture chambers coverglass II (Sarstedt). SiR-DNA (Spirochrome) was added for at least 5hrs prior to live-cell imaging. Images were acquired under a Zeiss AxioObserver Z1 epifluorescence microscopy system equipped with a heating and CO_2_ chamber (Digital Pixel) by using 40x/0.6 Plan-Neofluar or 40x/1.3 oil Plan-Apochromat objectives and a Hamamatsu ORCA-Flash4.0 LT Plus camera. For mitotic progression analysis, five to ten z-stacking images with 2μm intervals were taken with the indicated time intervals by using ZEN Blue software. Images were processed using ImageJ software and in-focus z-plane images were manually extracted to make image montages. For imaging of DNA thread formation in live cells, 40x/1.3 oil Plan-Apochromat objective was used to capture eight z-stack images with 800nm intervals and in-focus z-plane images were extracted using ImageJ software.

### Chromosome spread preparation

Following synchronisation using thymidine, cells were treated with pre-warmed hypertonic solution for 5–10 mins at 37°C (0.075M KCL). The swollen cells were then washed and fixed twice in methanol:acetic (3:1 ratio), before finally being re-suspended in fresh methanol:acetic solution. Chromosome spreads were dropped onto glass slides and either counterstained with Vectashield plus DAPI, or stored at room temperature for forthcoming FISH analyses. Colcemid was omitted in all mitotic spread preparations.

### Centromere & telomere Peptide Nucleic Acid (PNA) FISH

Centromere (CENPB-FAM; PNABio) & Telomere (Tel-Cy3; DAKO, Agilent) PNA probes were hybridised according to the manufacturer’s instructions. Briefly, chromosome spreads were rehydrated in TBS prior to fixation in 3.7% PFA solution. Slides were then washed and pre-treated before dehydration using a gradient ice-cold ethanol wash (70%, 90% and 100%). Slides were air dried and PNA probe added to spreads. Slides were then co-denatured at 80°C for 1 minute and incubated for 2 hours at room temperature, before counterstaining using DAPI Vectashield.

### Multi-colour FISH

mFISH was performed by using 24XCyte Human Multicolor FISH probe (MetaSystems) according to the manufacturer’s instructions. Images were acquired by MetaSystems using a Zeiss AxioObserver Z1 epifluorescence microscopy system with a CoolCube CCD camera and 100x/1.4 oil Plan-Aprochromat objective. mFISH karyotyping was carried out by using ISIS Imaging software.

### Immunoblotting

Cells were trypsinized and lysed on ice for 15–20 mins with lysis buffer (50mM Tris pH 7.5, 300mM NaCl, 5mM EDTA, 1% Triton X-100, 1.25 mM DTT, 1mM PMSF and cOmplete™ protease inhibitor cocktail). Protein concentration was quantified using a Bradford assay (Bio-Rad). Immunoblotting (IB) was performed following standard procedures. Primary antibodies used for IB in this study: anti-BLM (Abcam, ab2179, 1:2000), anti-PICH (Abnova; H00054821-B01P, 1:300), anti-GFP (Abcam, ab290, 1:1000) and anti-Ku80 (Abcam, ab80592, 1:10000).

### Flow cytometry

Cells were trypsinised, washed with PBS and fixed with 70% ice-cold ethanol. For cell cycle analysis, cells were washed with PBS and re-suspended in propidium iodide (PI) /RNaseA staining buffer. FACS profiles were then determined and analysed using BD Accuri C6 sampler.

### Kinetochore/centromere foci measurement

Samples were subjected to pre-extraction for 10–15sec followed by fixation and immunofluorescent staining as described above. Thirty to fifty z-stacking images with 200nm intervals were acquired and deconvolved using Huygens Professional deconvolution software (SVI). Kinetochore foci on each single z-plane were marked and measured using the ImageJ Plugins detailed below.

### ImageJ measurement of kinetochore foci coordinates, distances and intensities

#### Spot Pair Distance Tool

Measures the distance between spots in 2 channels of an image. The tool searches within a focus/box radius, typically +/−5px, for a local maxima in the two pre-selected analysis channels. The centre-of-mass around each maxima, typically +/− 2px, is computed as the centre of intensity for each channel. Dragging from the clicked point creates a reference direction. The Euclidean distance between the centres is reported, optionally with the signed XY distance and angle relative to the reference direction. Visual guides are overlaid on the image to assist in spot selection and direction orientation. Available in the latest GDSC ImageJ plugins.

#### Spot Fit Tool

Fits a 2D Gaussian to a spot in an image. The tool searches within a box radius, typically +/− 3px, for a local maxima in the pre-selected analysis channel. A 3×3 smoothing filter is applied before identification of the maxima. A 2D Gaussian function is then fitted to the data using non-linear least-squares fitting and poor fits rejected using a signal-to-noise ratio. The parameters for the fit are reported including the total intensity under the Gaussian function and the local background value. Visual guides are overlaid on the image to show the fitted location. Available in the pre-release GDSC SMLM ImageJ plugins.

### Statistics

Statistical analysis was performed using GraphPad Prism 7 software by two-tailed unpaired Student’s t-test and two-way ANOVA as per the experimental requirement.

## Supplementary Figure Legends

**Supplementary Figure 1.**
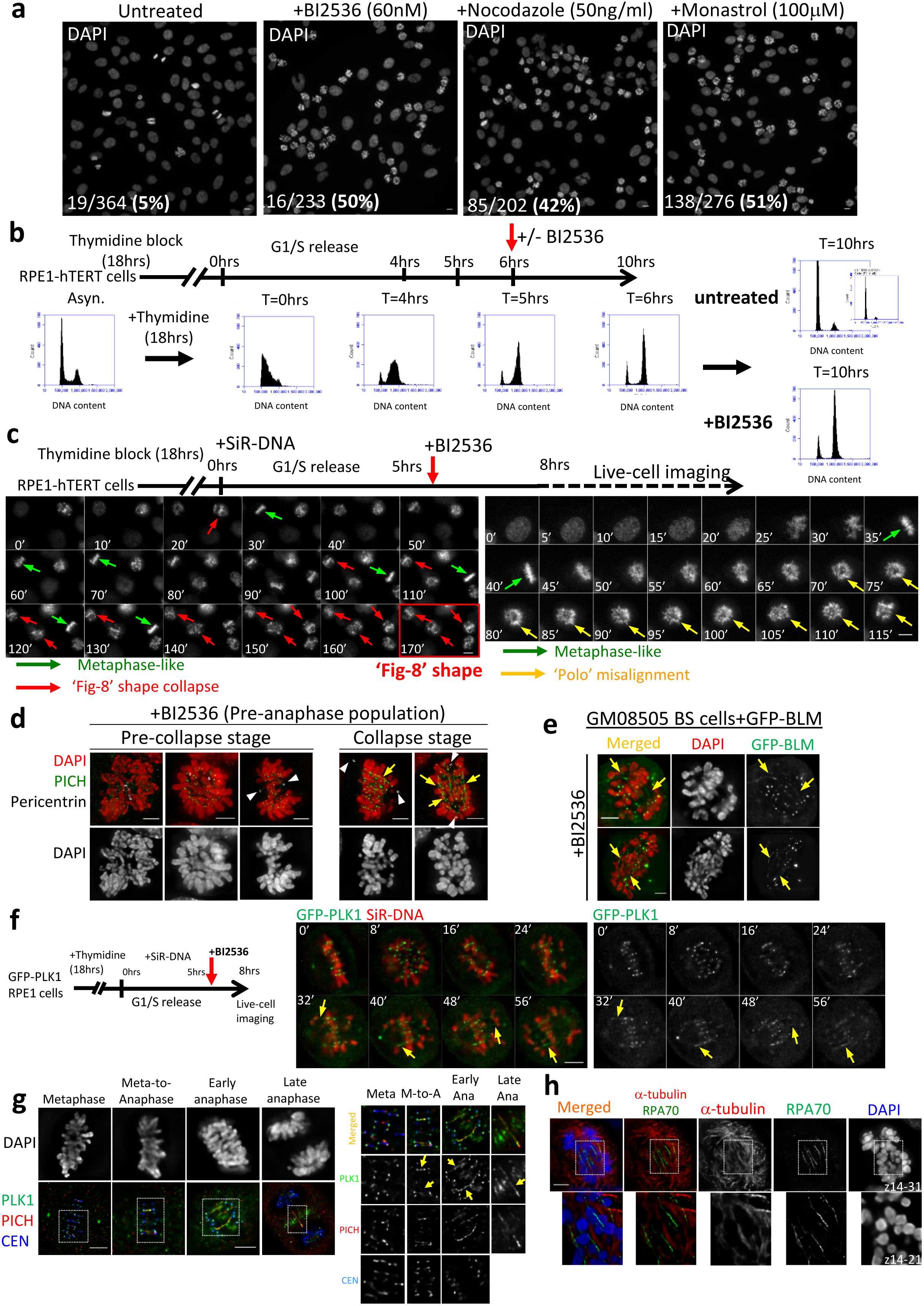
Plk1 inhibition by BI2536 small molecule inhibitor induces metaphase collapse and the formation of centromeric DNA threads. **a**, Asynchronous RPE1 cells were treated with or without the indicated inhibitors for 18hrs and stained with DAPI. Mitotic index is shown. Scale bar=10μm. **b**, FACS analysis of synchronised RPE1 cells with or without BI2536 treatment. Asynchronous RPE1 cells were treated with 2mM thymidine for 18hrs and released into fresh medium. BI2536 [60nM] was added at 6hrs post G1/S release. At indicated time points, samples were collected for FACS analysis. **c**, Time-lapse live-cell microscopy on RPE1 cells released from a single thymidine block. 60nM of BI2536 was added at 5hrs post G1/S release. Imaging started at 8hrs post release. SiR-DNA was used to label the DNA. Left panels showing an example of three cells entering mitosis and progressing into a metaphase(-like) stage (green arrows), followed by metaphase collapse with chromosome misalignment reminiscent of a ‘Figure-8’ shape (red arrows). Right panels show an RPE1 metaphase-like cell collapsing into a typical ‘Polo’ misalignment (yellow arrows). **d**, RPE1 cells were synchronised and treated with DMSO or BI2536 as (b) before being subjected to immunofluorescence staining using PICH and Pericentrin antibodies. Representative images of different mitotic populations before and after metaphase collapse (Arrows denote PICH-coated thread-like structures; arrowheads show centrosome positions). **e**, EGFP-tagged BLM protein binds to DNA threads (arrows) in Bloom’s syndrome fibroblasts following BI2536 treatment. **f**, Time-lapse live-cell imaging on pre-synchronised RPE1 cells stably expressing GFP-tagged PLK1 showed metaphase collapse and DNA thread formation (arrows) after BI2536 treatment. DNA was labelled using SiR-DNA. **g**, RPE1 cells were released from single thymidine block into fresh medium, followed by immunofluorescence staining after 8hrs. Deconvolved images showing ultrafine-DNA bridges (UFBs) (arrows) from metaphase to anaphase stained by PLK1, PICH and CEN antibodies. Enlarged regions are shown below. **h**, Representative images showing distinct localisation of DNA threads (labelled by RPA70) and microtubules (by α-tubulin) in BI2536-treated RPE1 cells. Scale bar=5μm.

**Supplementary Figure 2.**
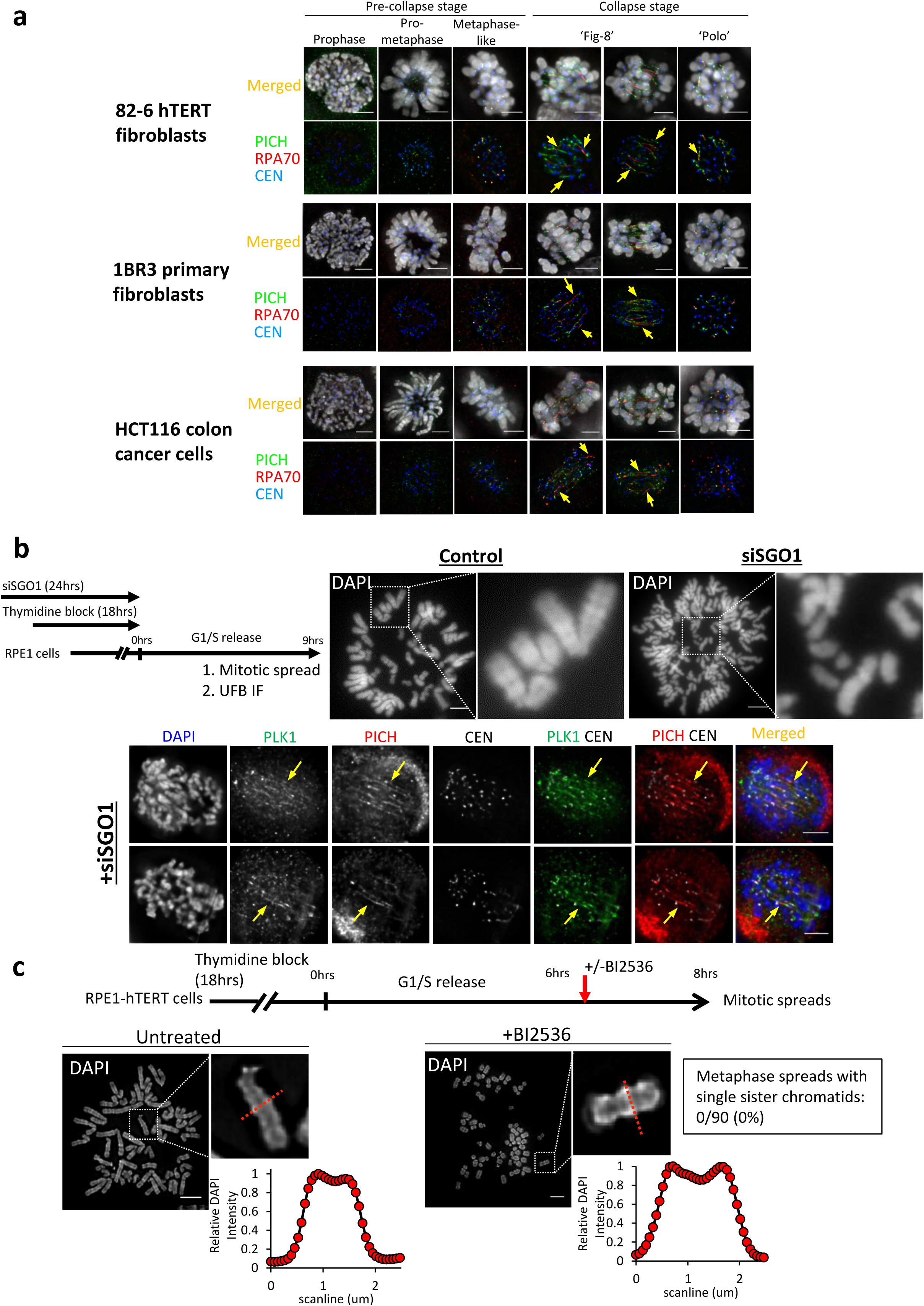
BI2536 induces <u>p</u>re-anaphase <u>i</u>nter-chromatin DNA <u>t</u>hread (PIT) formation in various human normal and tumour-derived cell lines, but does not induce premature loss of sister chromatid cohesion. **a**, Representative examples of different mitotic populations in pre-collapse and collapse stages showing PIT formation (arrows) in the indicated cell lines following thymidine synchronisation and BI2536 treatments. hTERT-immortalised 82–6 fibroblasts (upper), 1BR3 primary fibroblasts (middle) and HCT116 colon cancer cells (bottom panels). **b**, RPE1 cells were treated with siRNA oligos targeting Sgo1 during a thymidine block. Mitotic spreads and immunofluorescent staining were carried out at 9hrs post G1/S release to determine loss of sister chromatid cohesion and UFB formation (arrows). **c**, Mitotic spread analysis was carried out in RPE1 cells followed by short treatment of BI2536. Bottom: graphs showing the relative intensities of scanlines across sister chromatids. 90 mitotic spreads of BI2536-treated cells were analysed. DAPI was used to stain the mitotic spreads. Scale bar=5μm.

**Supplementary Figure 3.**
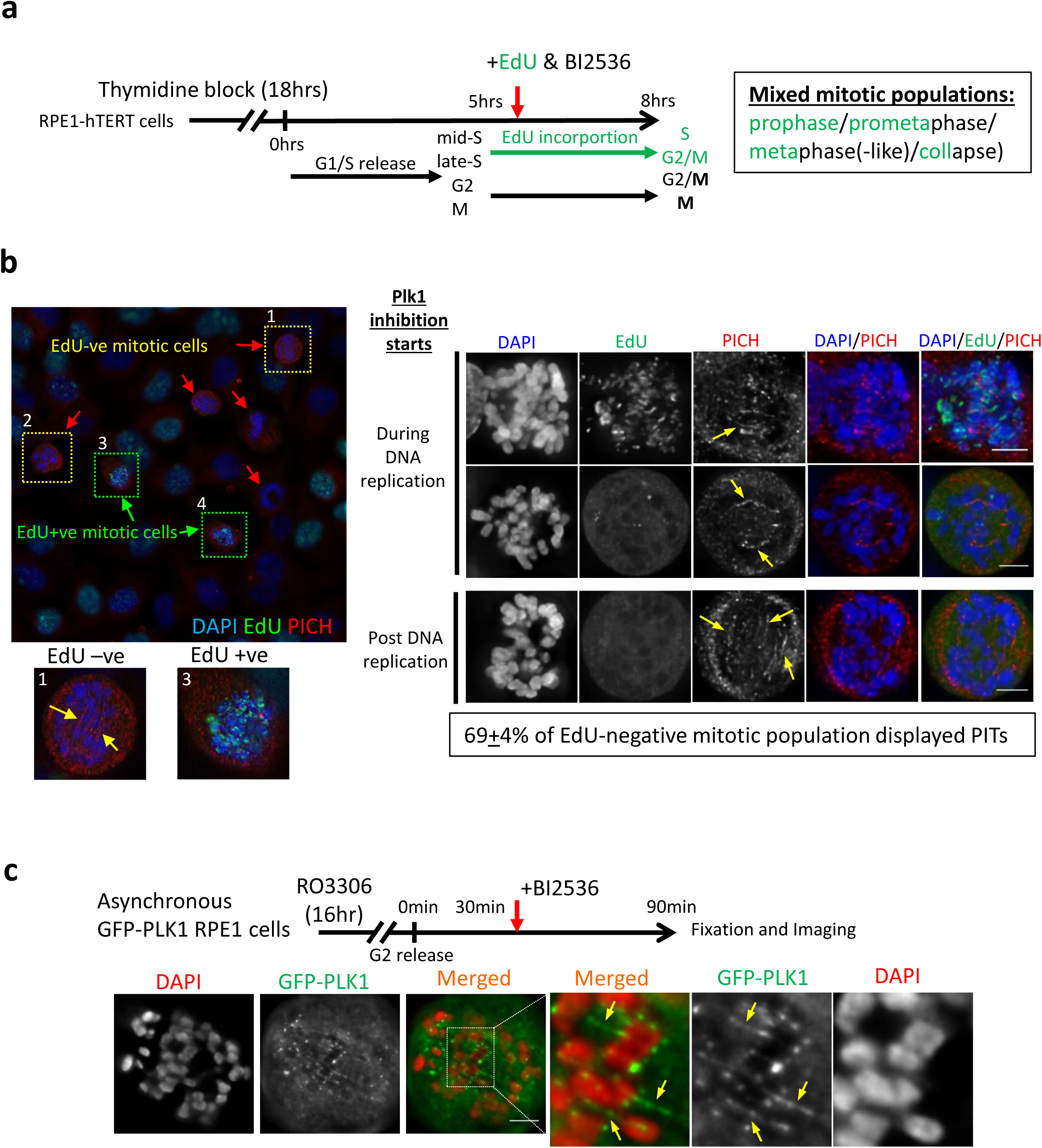
Constitutive mitotic Plk1 activity is required for the suppression of metaphase collapse and PIT formation. **a**, RPE1 cells were released from thymidine block. A thymidine analog, EdU, was added together with BI2536 at 5hrs post G1/S release to label on-going replicating cells. Immunofluorescence staining was performed at 8hrs post release. **b**, Left: An image showing a mixture of mitotic cells with or without EdU labelling signals (green). Right: Deconvolved images showing PIT formation (arrows) in EdU-positive and negative metaphase collapsed RPE1 cells. Percentage (mean ± S.D.) of BI2536-treated EdU-negative mitotic cells having PITs were scored in three independent experiments. **c**, Asynchronous GFP-tagged PLK1 RPE1 cells were arrested in G2 by the Cdk1 inhibitor, RO-3306, for 16hrs. 30min post G2 release, BI2536 was added for 60mins and then subjected to imaging. An example showing the formation of PITs (arrows) in metaphase collapsed GFP-PLK1 RPE1 cells. Scale bar=5μm.

**Supplementary Figure 4.**
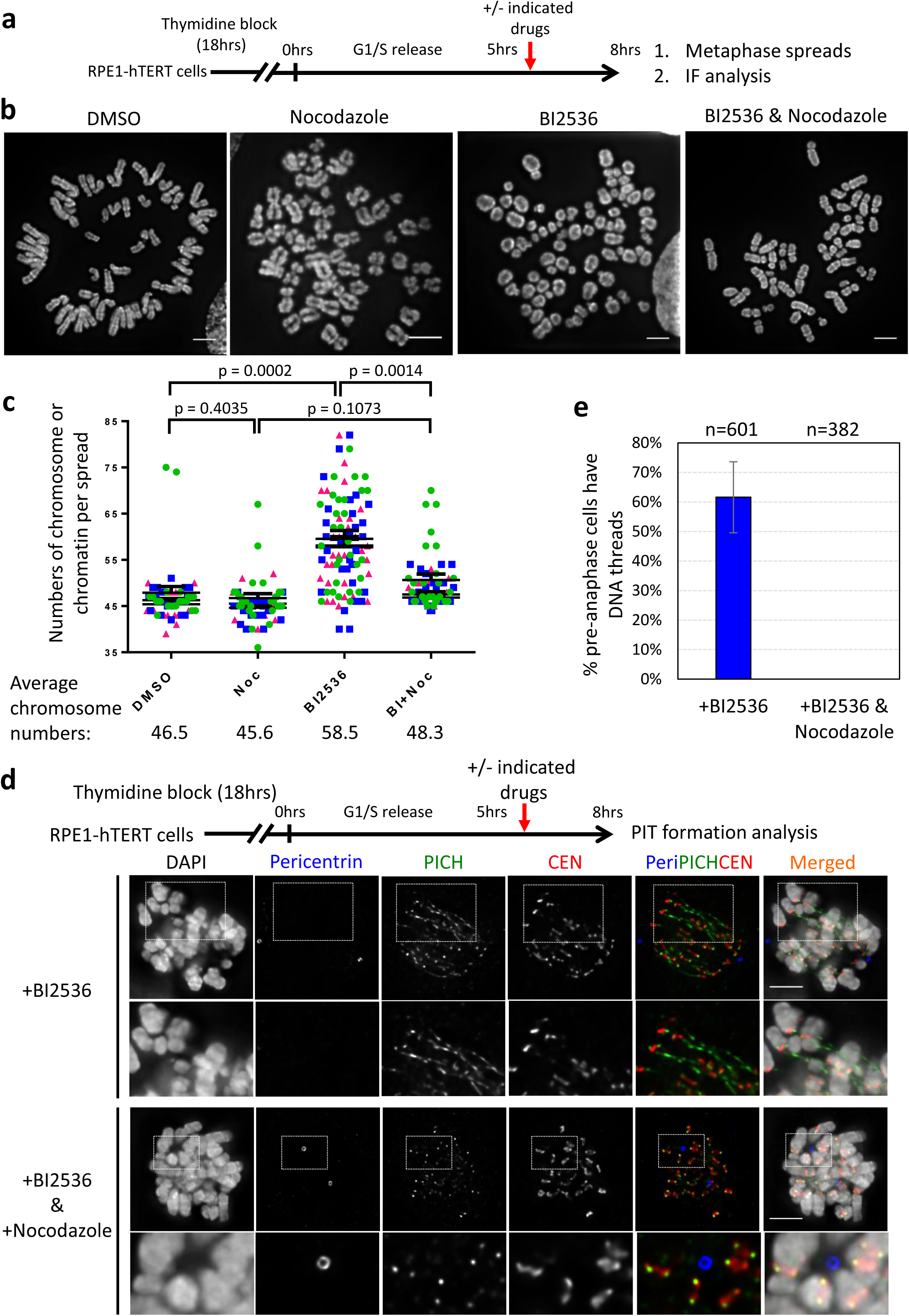
Spindle microtubules trigger chromosome fragmentation and PIT formation. **a**, Synchronisation and BI2536 treatment protocols on RPE1 cells. **b**, Representative examples of mitotic spread chromosomes after short treatments with the indicated inhibitors. **c**, Quantitation of chromosome/chromatin number in RPE1 cells treated as of (a) (A total of 90 spreads were scored from three independent experiments for each condition. Mean from each experiment is shown). **d**, Representative examples of mitotic arrested RPE1 cells showing PIT positive (BI2536) or negative (BI2536/NOC) formation after short treatments with the indicated inhibitors. **e**, Quantitation of PIT formation in RPE1 cells treated in (d), (n=total number of cells scored from three independent experiments. Mean±S.D. is shown). Scale bar=5μm.

**Supplementary Figure 5.**
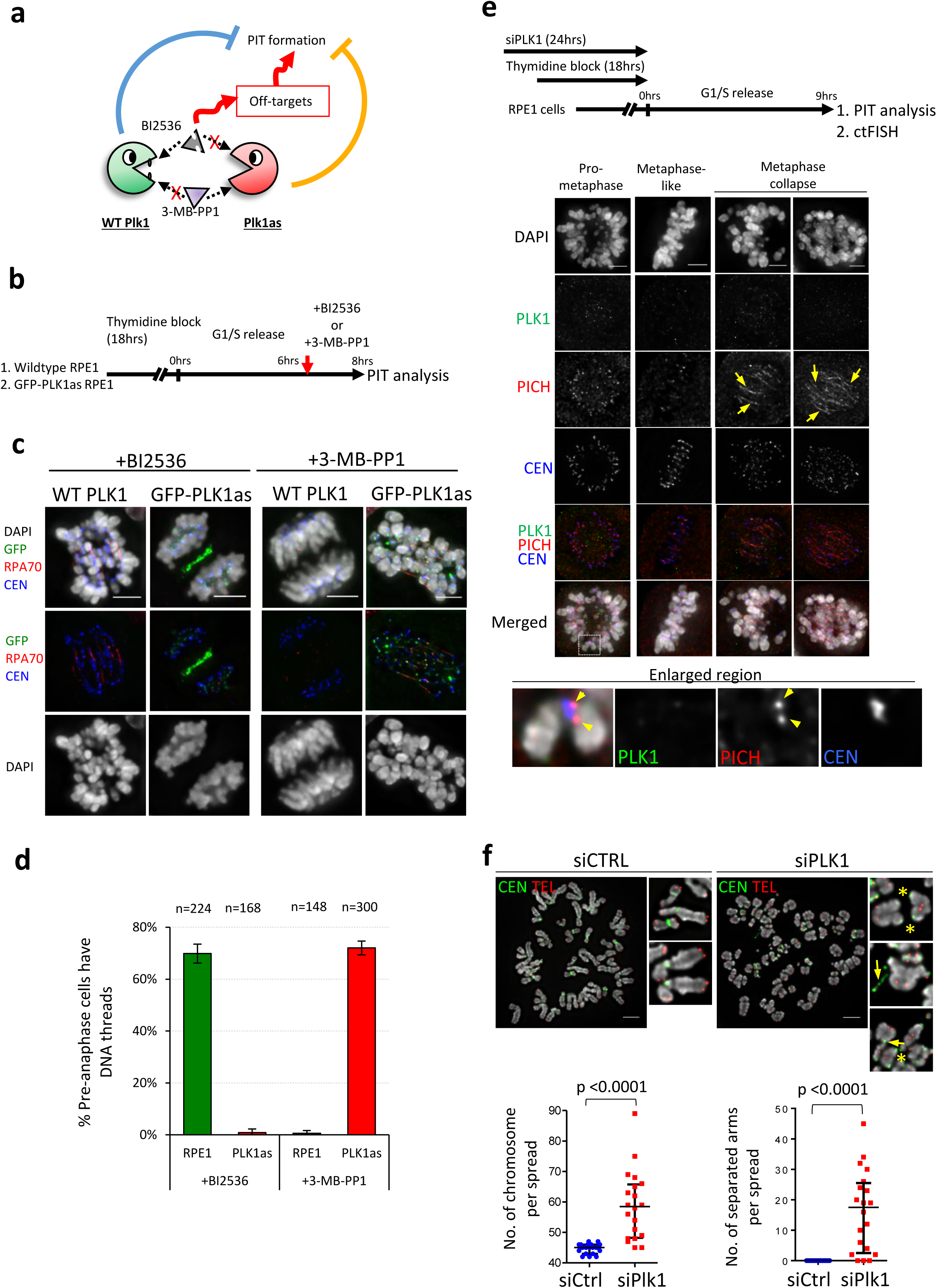
Plk1 kinase activity is required to suppress PIT formation. **a**, A diagram depicts the potential off-target effects on PIT formation by BI2536 small molecule, e.g. Plk2, Plk3 & Plk4 inhibition **b**, Wildtype RPE1 and GFP-tagged Plk1as RPE1 cells were synchronised by single thymidine block. Six hours post G1/S release, BI2536 [60nM] or 3-MB-PP1 analog [1μΜ] were added to cultures before PIT formation analysis. **c**, Examples of wildtype RPE1 and GFP-tagged Plk1as RPE1 mitotic cells showing PITs following BI2536 or 3-MB-PP1 treatments. Scale bar=5μm. **d**, Quantitation of PIT formation in wild-type and GFP-Plk1as RPE1 cells treated with either BI2536 or 3-MB-PP1 (mean±S.D. is shown; n=total numbers of cells scored across three independent experiments). **e**, RPE1 cells were treated with Plk1 siRNA oligos during thymidine arrest. Nine hours post G1/S release, cells were subjected to immunostaining and centromere-telomere FISH assays. Examples of RPE1 mitotic cells after Plk1 depletion showing PICH-coated PITs (arrows). Enlarged region: arrowheads denote PICH protein remains at kinetochores in the absence of PLK1. **f**, Examples of ctFISH on Plk1-knockdown RPE1 cells after release from thymidine block (asterisk, chromosome dislocation; arrows, centromeric DNA threads). Bottom, quantitation of chromosome/chromatin numbers and whole chromosome arm separation in the Plk1-depleted RPE1 cells, 19 siControl and 20 siPlk1 mitotic spreads were scored (mean±S.E.M).

**Supplementary Figure 6.**
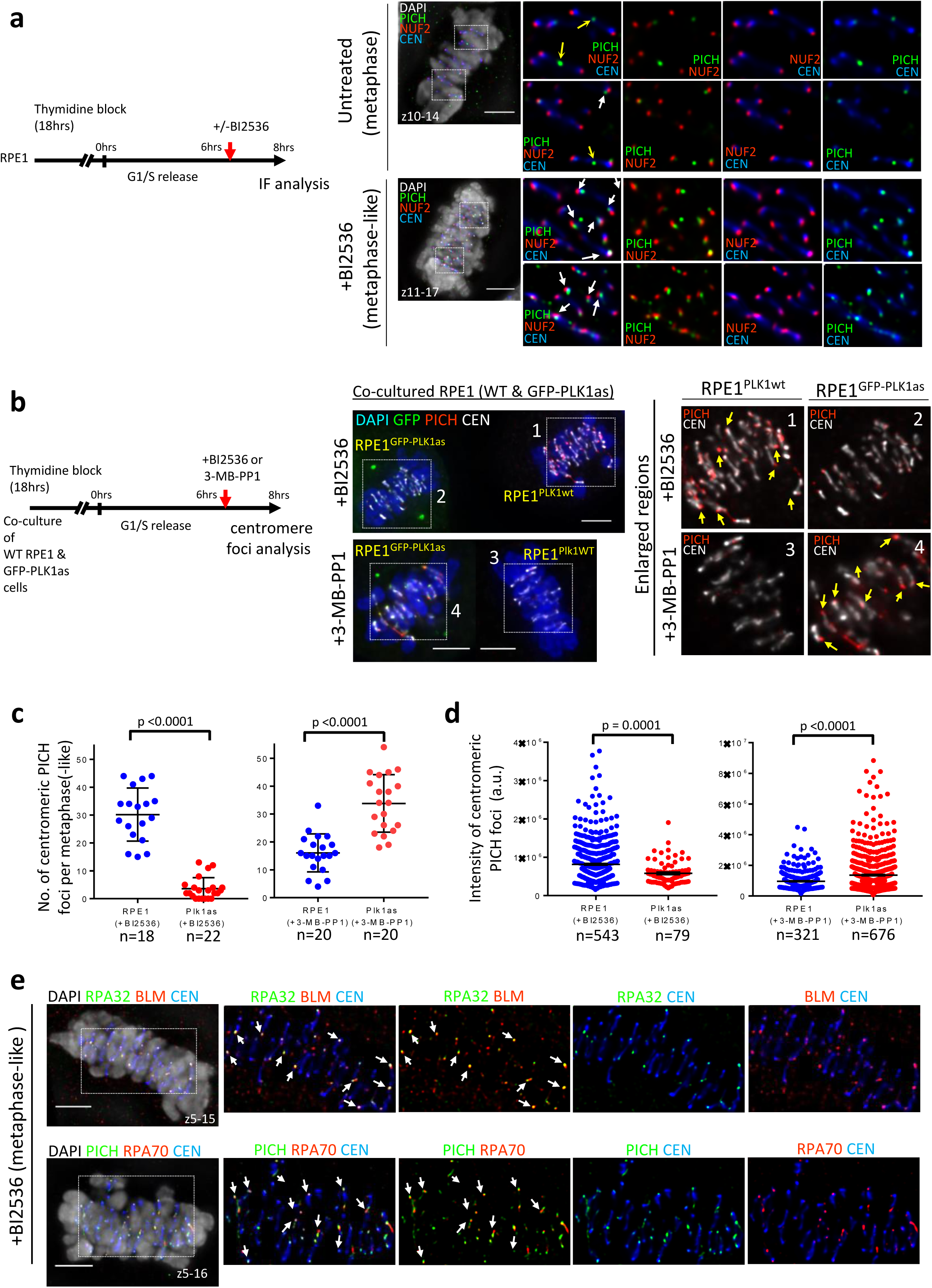
Accumulation of PICH, BLM and RPA at kinetochores in the pre-collapsed metaphase-like population. **a**, RPE1 cells were arrested by thymidine for 18hrs. BI2536 was added 6hrs post release and cells subjected to immunostaining with PICH, NUF2 and CEN antibodies. Examples of metaphase and metaphase(-like) cells were shown. NUF2 marks the kinetochore positions. Yellow arrows indicate PICH foci at the centre of centromeres; white arrows indicate PICH foci at/near kinetochores. **b**, Examples of co-culture of wildtype RPE1 and GFP-PLK1as RPE1 metaphase-like cells, stained with PICH, after treatments of either [60nM] BI2536 or [1μM] 3-MB-PP1 analog. Number **(c)** and intensities **(d)** of centromeric PICH foci measured from (b). n=total number of cells (c) and foci (d) scored; mean±S.D. (c) and mean+S.E.M. (d) are shown. **e**, Co-localisation of PICH, BLM and RPA proteins at kinetochores (arrows) in BI2536-treated metaphase(-like) RPE1 cells. Scale bar=5μm.

**Supplementary Figure 7.**
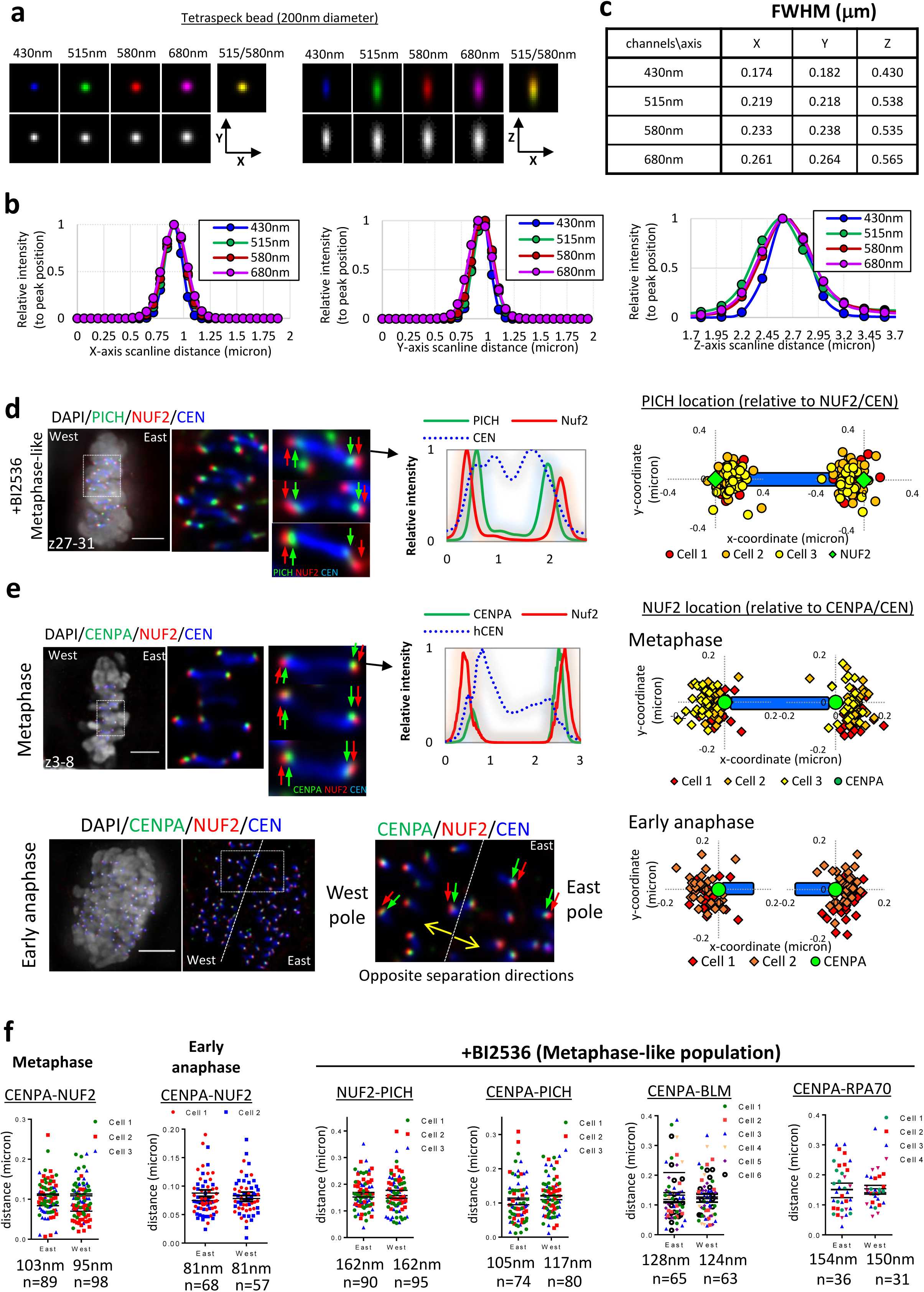
High resolution and precision deconvolution microscopy reveals UFB-binding complex underneath kinetochores. **a**, Images of a 4-color TetraSpeck bead of 200nm diameter in X, Y and Z. **b**, Scanlines showing the alignments of X, Y, Z positions of a TetraSpeck bead. **c**, Full width at half maximum (FWHM) of the deconvolved images of a TetraSpeck bead. **d**, Relative locations and distances of PICH to NUF2 in East and West poles of centromeres in BI2536-treated metaphase(-like) RPE1 cells. **e**, Relative locations and distances of NUF2 to CENPA in East and West poles of centromeres in normal RPE1 metaphase (upper panels) and early anaphase cells (bottom panels). Scale bar=5μm. **f**, Mean distances of inner and outer KT components as well as PICH/BLM/RPA proteins to kinetochores (left to right) (n=total numbers of foci analysed).

**Supplementary Figure 8.**
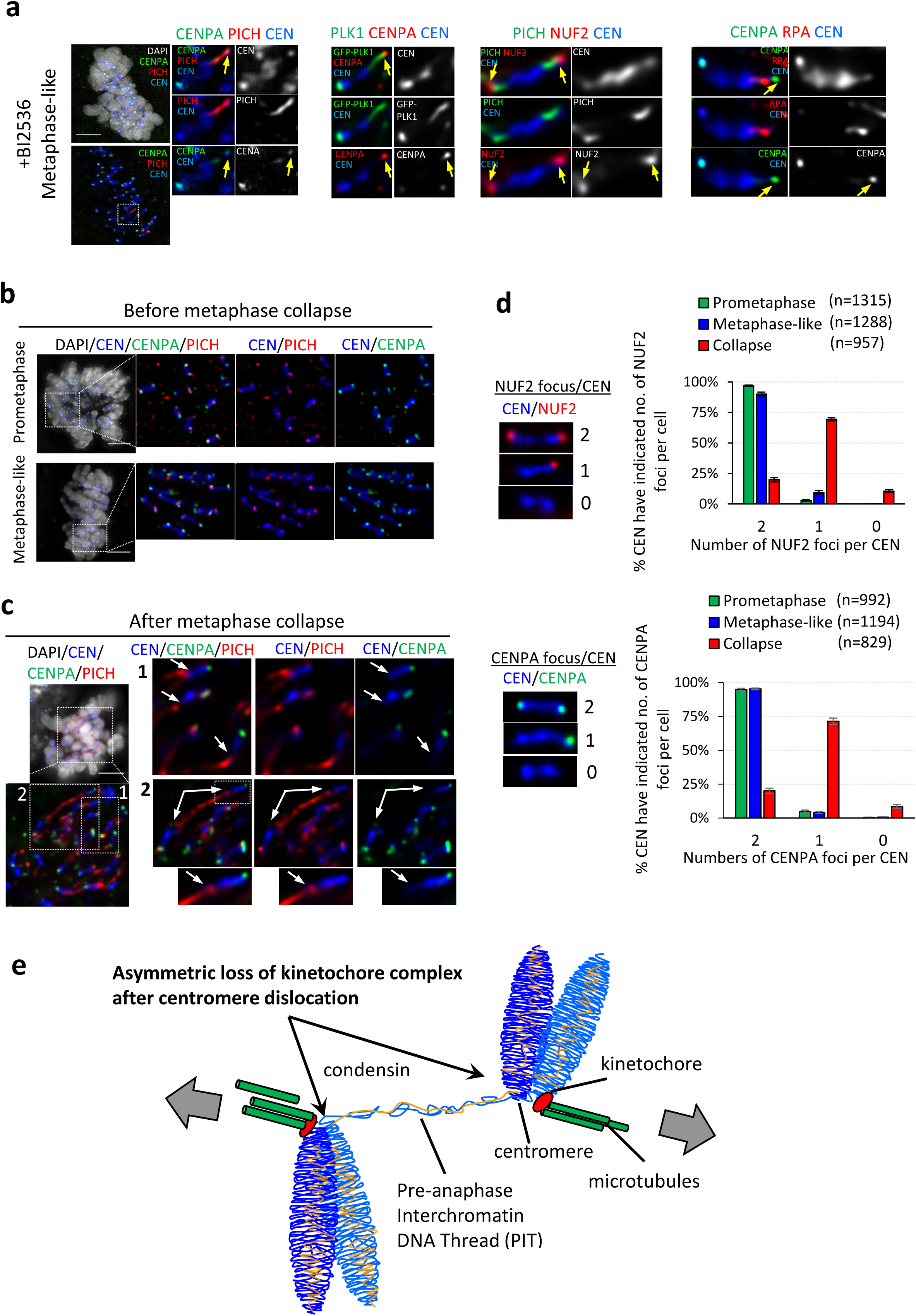
Detachment and loss of kinetochore complexes after Plk1 inhibition. **a**, Examples of centromeres displaying loosely attached kinetochores that remain connected by PIT threads. Representative images showing kinetochore foci at each centromere before **(b)** and after **(c)** metaphase collapse. **d**, Percentages of centromeres having the indicated numbers of kinetochores in prometaphase, metaphase(-like) and metaphase collapse populations after short treatments with BI2536. (n=total number of centromeres scored from 23 to 30 cells across three separate experiments, mean ± S.D. shown). Scale bar=5μm. **e**, A diagram depicts the formation of PITs and resultant centromere-kinetochore configuration after metaphase collapse induced by Plk1 inhibition. The split centromeres retain only one of the two kinetochores.

**Supplementary Figure 9.**
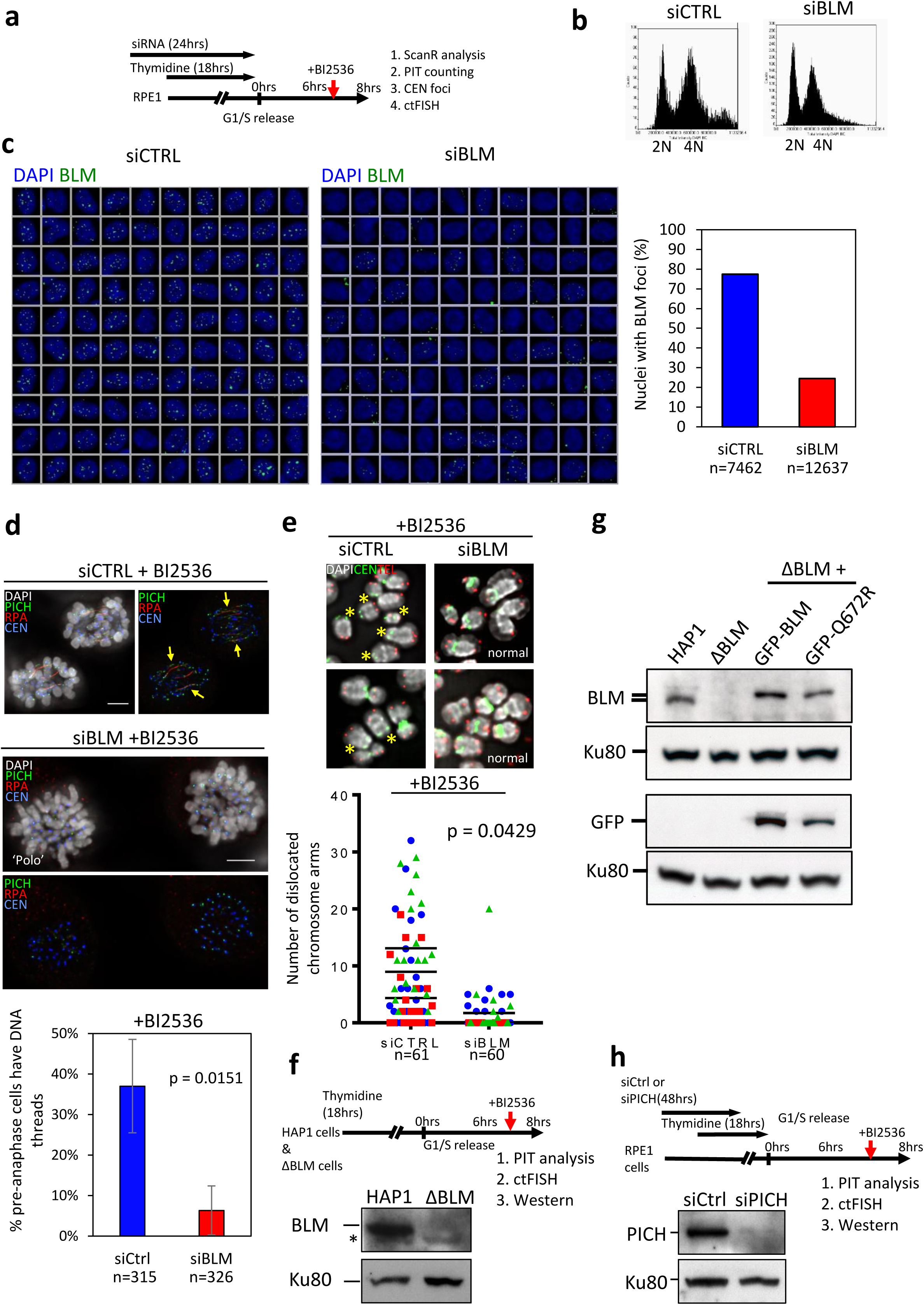
PICH and BLM trigger BI2536-induced centromere dislocation and PIT formation. **a**, RPE1 cells were treated with control or BLM siRNA oligos for 24hrs. Thymidine was added in the last 18hrs. Six hours post thymidine release, [60nM] BI2536 was added for 2 hours before fixation and IF analysis. **b**, Cell cycle profiles (measured by DAPI staining) of the siRNA transfected RPE1 cells after 8 hour release from thymidine block. **c**, Galleries of 100 random nuclei from (a) showing BLM foci. Right: percentages of RPE1 interphase nuclei showing spontaneous BLM foci. Slides were scanned and analysed by Olympus ScanR high-content screening station. **d**, Percentages of siControl and siBLM preanaphase populations showing PITs (mean ± S.D. from three independent experiments). **e**, Frequencies of centromere dislocation in siControl and siBLM RPE1 cells after BI2536 treatments (means from three independent experiment shown). n=total numbers of cells/mitotic spread analysed. **f**, Immunoblot of BLM knockout HAP1 cells (Asterisk, non-specific band). **g**, Immunoblot of BLM knockout HAP1 cells and its derivatives stably expressing a GFP-tagged wildtype BLM or Q672R mutant. **h**, Immunoblot of RPE1 cells after siControl and siPICH oligo transfection. Ku80 is used as loading control.

## Supplementary Movie Legends

**Movie S1. BI2536 induces metaphase collapse and mitotic arrest in normal RPE1 diploid cells.** RPE1 cells were released from single thymidine block and [60nM] BI2536 was added 5hrs post the release. Time-lapse live-cell imaging started at 8hrs post G1/S release with 10min intervals. DNA was stained using SiR-DNA.

**Movie S2. Inhibition of APC/C arrests cells in metaphase.** RPE1 cells stably expressing eGFP-PLK1 protein were released from single thymidine block and [12μM] ProTAME was added 6hrs post the release. Time-lapse live-cell imaging started at 8hrs post G1/S release with 10min intervals. DNA was stained using SiR-DNA.

**Movie S3. Inhibition of Plk1 in metaphase-arrested cells leads to metaphase collapse and PIT formation.** RPE1 cells stably expressing EGFP-PLK1 protein were released from single thymidine block and [12μM] ProTAME was added 6hrs post the release. At 8hrs post G1/S release, [60nM] BI2536 was added and time-lapse live-cell imaging started immediately. DNA was stained by SiR-DNA.

**Movie S4. Inhibition of Plk1 leads to chromosome misalignment following metaphase.** RPE1 cells were treated with control siRNA oligos for 48hrs. In the last 18hrs, thymidine was added to arrest cells in G1/S. [60nM] BI2536 was added 6hrs post the release. Time-lapse live-cell imaging started at 8hrs post G1/S release with 7min intervals. DNA was stained using SiR-DNA.

**Movie S5. Depletion of BLM delays metaphase collapse in BI2536-treated RPE1 cells.** RPE1 cells were treated with BLM siRNA oligos for 48hrs. In the last 18hrs, thymidine was added to arrest cells in G1/S. [60nM] BI2536 was added 6hrs post the release. Time-lapse live-cell imaging started at 8hrs post G1/S release with 7min intervals. DNA was stained using SiR-DNA.

**Movie S6. Depletion of PICH delays metaphase collapse in BI2536-treated RPE1 cells.** RPE1 cells were treated with PICH siRNA oligos for 48hrs. In the last 18hrs, thymidine was added to arrest cells in G1/S. [60nM] BI2536 was added 6hrs post the release. Time-lapse live-cell imaging started at 8hrs post G1/S release with 7min intervals. DNA was stained using SiR-DNA.

